# Molecular Forecasting of Domoic Acid during a Pervasive Toxic Diatom Bloom

**DOI:** 10.1101/2023.11.02.565333

**Authors:** John K. Brunson, Monica Thukral, John P. Ryan, Clarissa R. Anderson, Bethany C. Kolody, Chase James, Francisco P. Chavez, Chui Pin Leaw, Ariel J. Rabines, Pratap Venepally, Hong Zheng, Raphael M. Kudela, G. Jason Smith, Bradley S. Moore, Andrew E. Allen

**Author notes:** These authors contributed equally to this work (John K. Brunson, Monica Thukral).

## Abstract

In 2015, the largest recorded harmful algal bloom (HAB) occurred in the Northeast Pacific, causing nearly 100 million dollars in damages to fisheries and killing many protected marine mammals. Dominated by the toxic diatom *Pseudo-nitzschia australis*, this bloom produced high levels of the neurotoxin domoic acid (DA). Through molecular and transcriptional characterization of 52 near-weekly phytoplankton net-tow samples collected at a bloom hotspot in Monterey Bay, California, we identified active transcription of known DA biosynthesis (*dab*) genes from the three identified toxigenic species, including *P. australis* as the primary origin of toxicity. Elevated expression of silicon transporters (*sit1*) during the bloom supports the previously hypothesized role of dissolved silica (Si) exhaustion in contributing to bloom physiology and toxicity. We find that co-expression of the *dabA* and *sit1* genes serves as a robust predictor of DA one week in advance, potentially enabling the forecasting of DA-producing HABs. We additionally present evidence that low levels of iron could have co-limited the diatom population along with low Si. Iron limitation represents a previously unrecognized driver of both toxin production and ecological success of the low iron adapted *Pseudo-nitzschia* genus during the 2015 bloom, and increasing pervasiveness of iron limitation may fuel the escalating magnitude and frequency of toxic *Pseudo-nitzschia* blooms globally. Our results advance understanding of bloom physiology underlying toxin production, bloom prediction, and the impact of global change on toxic blooms.

**Significance:** *Pseudo-nitzschia* diatoms form oceanic harmful algal blooms that threaten human health through production of the neurotoxin domoic acid (DA). DA biosynthetic gene expression is hypothesized to control DA production in the environment, yet what regulates expression of these genes is yet to be discovered. In this study, we uncovered expression of DA biosynthesis genes by multiple toxigenic *Pseudo-nitzschia* species during an economically impactful bloom along the North American West Coast, and identified genes that predict DA in advance of its production. We discovered that iron and silica co-limitation restrained the bloom and likely promoted toxin production. This work suggests that increasing iron limitation due to global change may play a previously unrecognized role in driving bloom frequency and toxicity.

## Introduction

Harmful algal blooms (HABs) produce a variety of potent molecular toxins, including many that pose an active threat to human health (1). Bioaccumulation of the diatom-produced neurotoxin domoic acid (DA) in various seafoods such as Dungeness crab and mussels routinely triggers fishery closures around the world, resulting in millions of dollars of economic losses during toxic events (2). Acute human exposure to DA can lead to amnesic shellfish poisoning (ASP), a potentially lethal syndrome characterized by seizures, vomiting and short-term memory loss (3, 4). Because of the risks to human health afforded by DA and other algal toxins, routine environmental monitoring is implemented to better predict and respond to major HAB events (5–7).

Blooms of the DA-producing diatom genus *Pseudo-nitzschia* are near-annual events along the North American West Coast and are fueled by strong seasonal upwelling in the California Current Ecosystem (CCE) (8, 9). In 2015, a bloom of *Pseudo-nitzschia australis* (*P. australis*) extended from the Gulf of Alaska to Point Conception, California from spring to late summer and, to date, is the largest and longest recorded HAB in the Northeast Pacific Ocean. Toxic populations of *P. australis*, a species common to the central California coast, expanded their range northward in association with an extreme marine heatwave in 2014-2015 (10, 11). During this historically large HAB event, exceptionally high levels of DA were detected in Monterey Bay, California, prompting further study into the oceanographic mechanisms fueling the bloom event in the bay (12, 13). Monterey Bay experiences frequent phytoplankton blooms driven by strong seasonal upwelling at Point Año Nuevo to the north and Point Sur to the south (Fig. S1) (14–16). Such upwelling waters, with elevated concentrations of dissolved nitrogen and phosphorous can often drive phytoplankton communities into Si and/or iron limitation (17–19). Iron limitation has been shown to limit growth of coastal planktonic communities, and iron stress has been shown to upregulate DA production (20–22). The highly toxic *P. australis* HAB of 2015 coincided with historically low Si concentrations recorded throughout Monterey Bay, and limitation of this key nutrient has been shown to induce DA production in laboratory culture and modeling experiments (12, 23–27). However, a wide variety of biotic and abiotic factors have been shown to influence toxin production by *Pseudo-nitzschia*. Despite decades of research on DA production in the lab and in mesoscale oceanographic studies, a predictable set of physiological mechanisms underpinning DA production in the environment remains unclear.

The DA biosynthetic pathway in *Pseudo-nitzschia multiseries* was first discovered by using an RNA-sequencing approach to identify DA biosynthetic (*dab)* genes that were upregulated under simultaneous phosphate limitation and elevated *p*CO_2_, conditions that increase DA production in laboratory cultures (28, 29). The enzymes encoded by *dabA*, *dabC* and *dabD* were demonstrated to perform key biosynthetic reactions required for *in vitro* production of the DA molecule (28, 30). Among diatoms, *dab* genes have only been described within the genus *Pseudo-nitzschia* in the species *P. multiseries*, *P. multistriata*, *P. australis*, and *P. seriata* (28, 31). Following this initial discovery, homologues to the *dab* genes have also been characterized in red algae that produce either DA or the structurally related neurochemical, kainic acid (32, 33).

Molecular methodology, targeting DNA and/or RNA, can enable rapid identification of HAB species at significantly improved precision, and has been successfully deployed for routine monitoring of *Pseudo-nitzschia* in Monterey Bay (34). Additionally, detection of toxin biosynthetic genes has been proposed as a possible route for monitoring and forecasting HAB toxicity, and the cyanobacterial HAB community has implemented molecular monitoring guided by the biosynthetic genes for toxins such as microcystin and saxitoxin (35, 36). Environmental detection of *dab* transcripts is currently not implemented in the monitoring of DA-producing *Pseudo-nitzschia* HABs, and at present it is unknown how *dab* transcription might be related to DA production in the marine environment.

In this study, we targeted rRNA and mRNA to combine 18S and ITS2 metabarcoding with polyA-enriched RNA-sequencing to provide a framework for characterizing toxic *Pseudo-nitzschia* species, *dab*, and other gene transcription at the molecular level. These new metabarcoding and metatranscriptomic resources were generated from 52 near-weekly phytoplankton net-tow samples collected at Monterey Municipal Wharf II (MWII) in Monterey Bay, California from late 2014 to the end of 2015, encapsulating the entire *P. australis* bloom event (Fig. 1). Analysis of molecular barcoding datasets revealed the phytoplankton succession as community composition transitions from non-toxic diatom species to a near-monospecific bloom of DA-producing *Pseudo-nitzschia australis*. We also measured transcription of *dab* genes in the metatranscriptomics dataset and compared these findings with DA measurements taken at MWII and abundances of toxigenic species as revealed by metabarcoding. We focused on how expression patterns of *dab* genes and DA production related to a silicon transporter (*sit1*), a carbonate-dependent phytotransferrin, iron starvation-induced protein (*ISIP2A*), and ferritin (*FTN*). In addition, the metatranscriptomics data allowed us to further investigate highly expressed transcripts to describe the shifting physiology of the *P. australis* bloom throughout the entire HAB event.

**Figure 1.**
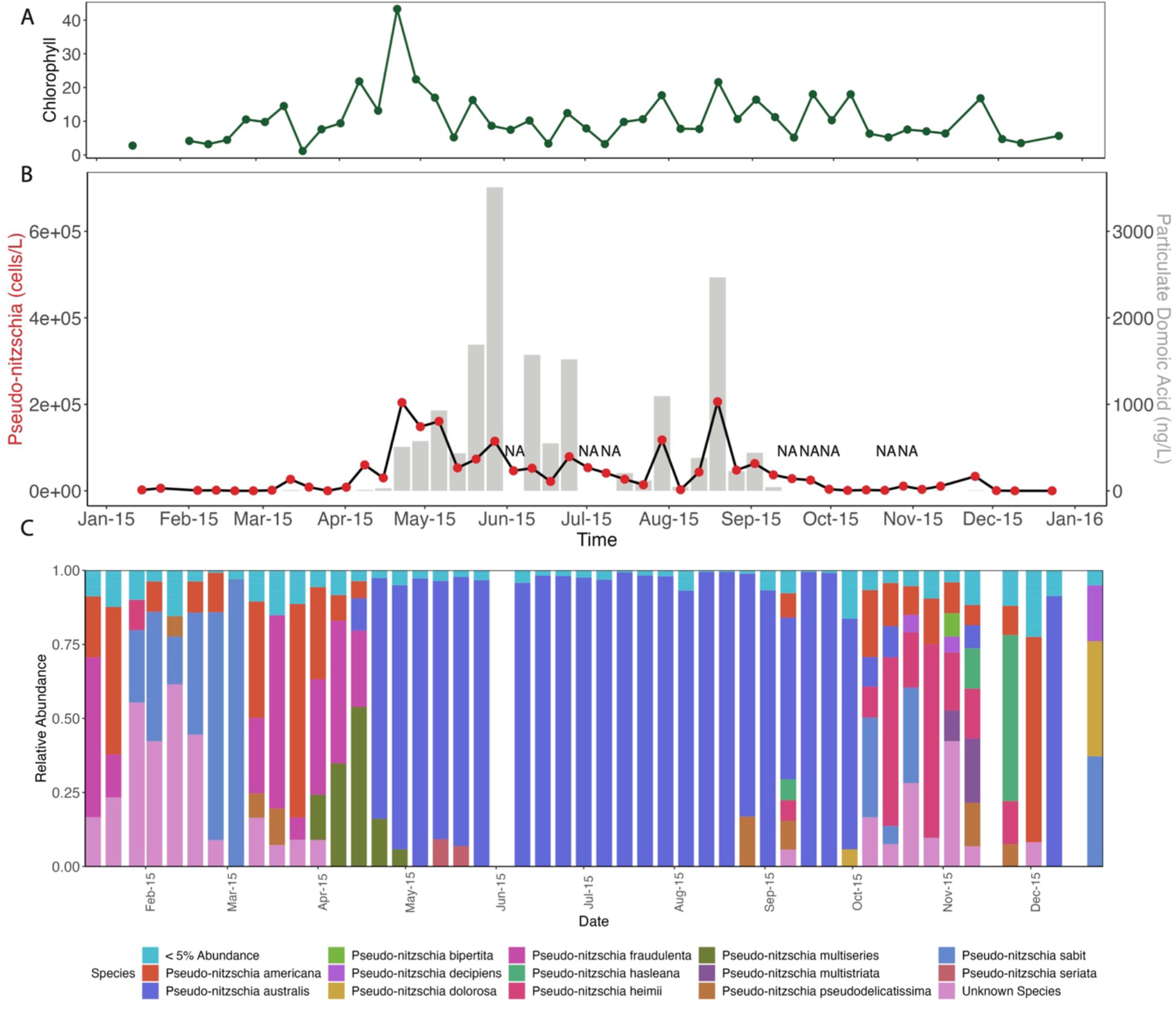
*Pseudo-nitzschia* Targeted ITS2 Amplicon Sequencing. (A) Chlorophyll*-a* (mg/m^3^) at MWII, (B) *Pseudo-nitzschia* cell counts from weekly MWII phytoplankton net tows (red dots, left axis) overlayed with particulate domoic acid (pDA, ng/L) measurements from filtered seawater (gray bars, right axis), as reported by CalHABMAP monitoring^7^ and (B) Relative abundance of *Pseudo-nitzschia* species as determined by ITS2 amplicon sequencing. Non-ITS2 sequences were removed prior to analysis. NA indicates missing data.

## Results and Discussion

### Amplicon Sequencing Reveals the Spring Phytoplankton Succession

Temperature and nitrate measurements at MWII in 2015 suggested that cold, nutrient-rich upwelling began in late March and early April, supported by geophysical and ecosystem dynamics studies (Fig. S2) (12). Amplicon sequencing of the 18S rRNA-V4 region revealed a stark transition from a zooplankton-dominant ecosystem characterized by the abundant dinoflagellate (Myzozoa) to a diatom-dominant (Bacillariophyta) community in early to mid-March, a shift coincident with increased chlorophyll concentrations at MWII (Fig. S3A, Fig. 1A). Diatoms remained the dominant members of the microbial community until mid-July when their relative abundance briefly decreased. The diatom-dominated phytoplankton bloom returned once more in August before its ultimate demise at the end of September, giving way to an intermittent assemblage of diatoms and dinoflagellates, copepods, and other grazing zooplankton in the Autumn months.

The early spring bloom was dominated by centric diatoms, primarily *Chaetoceros*, prior to transitioning in late-April to a community composed almost entirely of *Pseudo-nitzschia australis* that dominated the rest of the bloom season through early autumn (Fig. S3B, Fig. S4, Fig. S5). Such phytoplankton succession from a mixed assemblage community during new upwelling to a diatom-dominated community post-upwelling has been previously reported for the CCE (37, 38). These observations are further supported by chloroplast sequences recovered from 16S rRNA sequencing (Fig. S6). Returning diatom assemblages present in October and November were, like the early spring phytoplankton community, primarily composed of *Chaetoceros* (Fig. S6). Alpha-diversity analysis of the 18S-V4 data revealed that diatom diversity indeed declined during the bloom event, concurrent with previous observations suggesting a near-monospecific bloom of *P. australis*. This significant reduction in diversity was primarily driven by pennate diatoms resulting from *Pseudo-nitzschia* dominance, with centric diatom diversity appearing largely unaffected throughout the *P. australis* bloom (Fig. S7A & Fig. S7B).

### ITS2 Sequencing reveals *Pseudo-nitzschia* species diversity prior to *P. australis* dominance

While the Spring phytoplankton succession was resolved to the genus level by 18S rRNA metabarcoding, amplicon sequencing of the internal transcribed spacer 2 (ITS2) region was implemented to improve characterization of *Pseudo-nitzschia* species succession. The ITS2 region has been demonstrated to provide species-level resolution consistent with classical methods of species delineation, such as electron microscopy (39). ITS2 metabarcoding confirmed *P. australis* as the dominant species during the bloom event (10, 13). The early spring phytoplankton succession dominated by *Chaetoceros* appears to also include low levels of various *Pseudo-nitzschia* species such as *P. fraudulenta, P. cuspidata*, *P. americana*, and *P. pungens*. All these species, barring *P. americana*, have been demonstrated to produce low-to-modest amounts of DA in the lab, and their presence here coincides with very low levels of DA production detected at MWII (∼7 ng/L, March 11^th^, Fig. 1) (40, 41).

Beginning in late April, the community shifted to near-complete dominance of *P. australis*, the species previously identified to be the main driver of the bloom in Monterey Bay and along the entire North American West Coast (10, 12, 13). In addition, a substantial subpopulation of potentially toxic *P. multiseries* was also identified throughout the month of April in conjunction with modestly increased, but still low, DA concentrations (∼22 ng/L, April 15^th^, Fig. 1). Detection of the *P. multiseries* subpopulation at MWII is corroborated by *P. multiseries* presence during a similar period at the northern point of Monterey Bay at Santa Cruz Wharf (13). A third subpopulation of the potentially highly toxigenic *P. seriata* was also observed in early to mid-May, albeit at a much lower level of abundance than *P. australis*. These data suggest that several species may have contributed to DA production throughout different phases of the bloom season, and that a sizable population of *P. multiseries* may have been a significant contributor to initial DA production at MWII prior to the emergence of *P. australis* as the dominant species.

### Expression of *dab* genes reflects toxin production by *Pseudo-nitzschia*

We reconstructed the sequence and expression profiles of several *dab* transcripts in the polyA-enriched RNA-sequencing dataset. These transcripts displayed nearly identical nucleotide sequences compared to known *dab* genes from *P. australis, P. multiseries*, and *P. seriata*, three species identified by ITS2 sequencing during the bloom (Table S1) (28, 31). The full suite of core DA biosynthetic genes (*dabA*, *dabC*, and *dabD*) as expressed by the dominant bloom species, *P. australis*, was identified and were detected throughout the entire bloom event (April 15^th^ – September 30^th^). Assemblies also yielded *P. multiseries* transcripts for *dabA* and *dabC*, both of which displayed expression profiles coinciding with the detection of the species through ITS2 amplicon sequencing (April 1^st^ – mid May). Two additional *dab* genes were identified from the *de novo* assembly: one could be confidently assigned to *P. seriata* (*dabA*, April 22^nd^ – June 10^th^), and the other is of unknown origin (*dabC*, March 11^th^ - April 8^th^). The unknown *dabC* transcript was present early in the spring phytoplankton succession, coincident with the presence of various low-toxicity *Pseudo-nitzschia* spp. and low-level detection of particulate domoic acid (pDA). Active *dabA* and *dabC* gene transcription co-occurred with detection of pDA throughout most of the spring and summer months (Table S2, Fig. S8).

Phytoplankton sampling and pDA measurements taken from MWII, combined with molecular barcoding and *dab* transcription measurements, suggested two phases to the persistent *P. australis* bloom as observed from MWII. The first phase involved proliferation of the bloom and steady build-up of cellular DA (cDA), or pDA per *Pseudo-nitzschia* cell, throughout April and May in conjunction with *dab* gene expression by the *P. australis* and *P. multiseries* subpopulations during these months (Fig. S8A). The second phase of the bloom occurred after the July disappearance of *Pseudo-nitzschia* and subsequent decrease in cDA (Fig. 1, Fig. S8B). Beginning in late July with the sudden resurgence of *Pseudo-nitzschia* cells and DA, periodic increases in cDA were accompanied by increases in the abundance of *P. australis dabA* and *dabC* transcripts during the late phase of the *Pseudo-nitzschia* HAB (Fig. 1B, Fig. S8A). In both early and late phases of the bloom, increases in cDA concentrations coincided with spikes in transcript abundance for *dabA* and *dabC*, the two potentially diagnostic DA biosynthetic genes (Fig. S8B-D) (28).

### Expression profiles of *P. australis* transcripts reveal evolution of the bloom

Weighted gene correlation network analysis (WGCNA) on a highly-expressed subset of the *de-novo* assembled *P. australis* HAB metatranscriptome identified seven modules of transcripts with similar expression profiles (Fig. 2, Fig. S9) (42). Both *dabA* and *dabD* were found in module 7, along with two genes encoding key isoprenoid biosynthesis enzymes: 4-hydroxy-3-methylbut-2-enyl-diphosphate synthase (HDS) and isopentenyl-diphosphate delta-isomerase (IDI) (Fig. S10A). Both proteins are directly implicated in biosynthesis of the DA precursor geranyl pyrophosphate and are known to be co-expressed with the *dab* gene products (31). Two additional isoprenoid biosynthesis transcripts, 1-deoxy-D-xylulose-5-phosphate synthase (DXS) and 4-hydroxy-3-methylbut-2-enyl diphosphate reductase (HDR), were found in modules 2 and 4, respectively. All four isoprenoid biosynthesis genes included in this analysis are implicated in the chloroplastic, non-mevalonate pathway thought to feed directly into DA biosynthesis (28, 31, 43).

**Figure 2.**
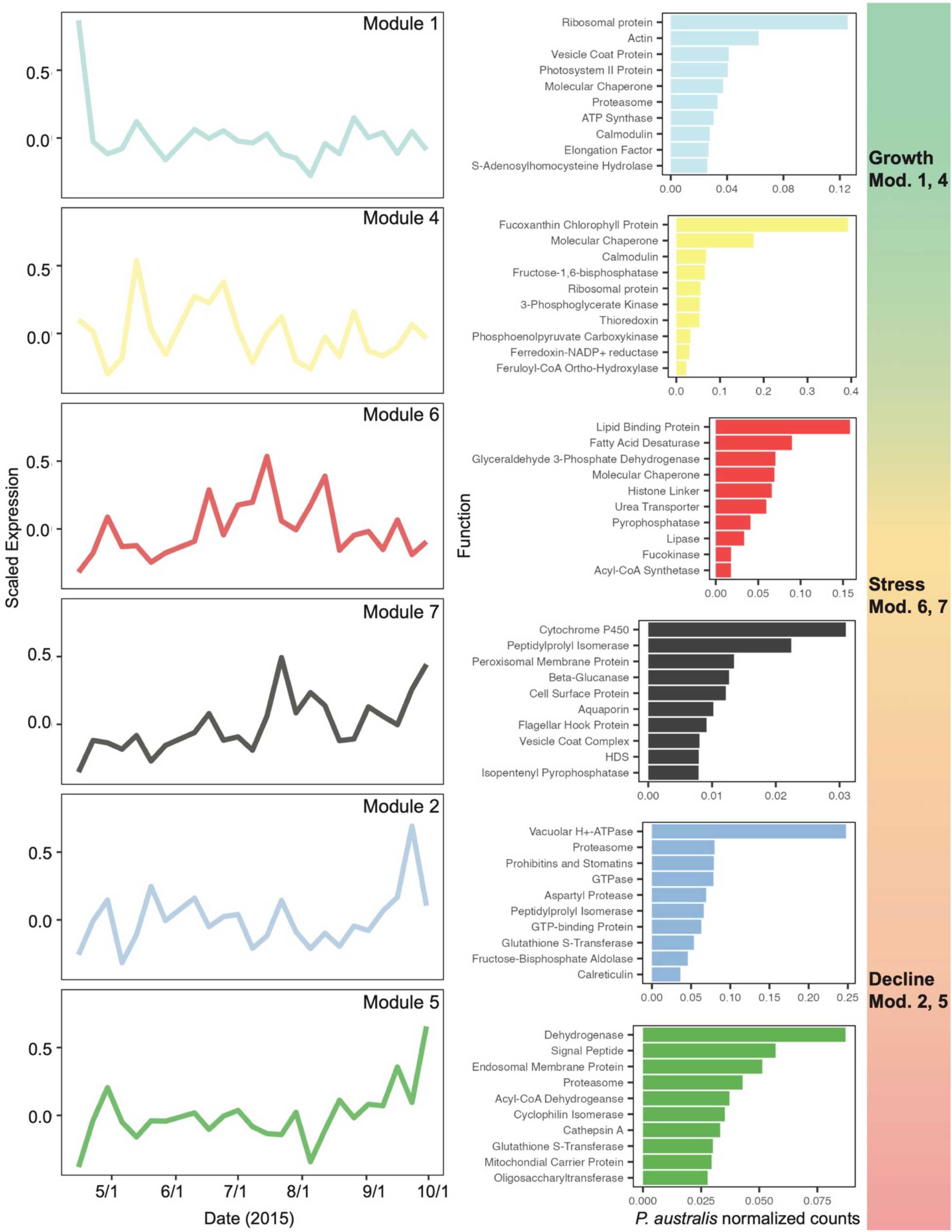
Patterns in co-expressed *P. australis* genes reveal bloom progression from growth to stress to decline during the duration of the HAB. The data represent relative expression modules for *P. australis* transcripts throughout the HAB as determined by WGCNA analysis. Corresponding bar charts on the right of each module show the ten most abundant functions of open reading frames (ORFs) of each module using KEGG descriptions. The data is shown as the sum of *Pseudo-nitzschia australis* normalized read counts. ORFs without annotations are not shown.

The remaining DA biosynthetic gene *dabC* was found in module 4, together with HDR but separate from *dabA* and *dabD* (Fig. S10A). While *dabC* is co-expressed with *dabA* and *dabD* in *P. multiseries* and *P. seriata* cultures, its enzyme DabC catalyzes the third biosynthetic reaction to DA in a suspected different cellular compartment than the DabA and DabD enzymes and thus might be differentially regulated (28, 31). Module 4 also displayed functional clustering of photosynthetic machinery like chlorophyll binding proteins, light harvesting complexes, the iron-sulfur subunit of the cytochrome b6f complex, photosystem I reaction center subunit VIII, photosystem II stabilizing-protein PsbU, ferredoxin-NADP^+^ reductase, transketolase, and fructose-1,6-bisphosphatase (44) (Fig. S10B).

Module 1 includes transcripts relatively abundant at the beginning of the bloom and are among the first *P. australis* transcripts detectable. Many of these transcripts are related to active cell growth and proliferation. Half of the 16 ribosomal subunit transcripts in the WGCNA dataset were found in module 1 (Fig. S10B). Five ribosomal subunits included in the dataset are predicted to reside in the chloroplast, all of which were found in module 1, potentially indicating increased chloroplast ribosome biogenesis early in the bloom progression. Transcripts for F-type ATP synthase subunits were also primarily found in module 1, suggesting increased ATP production in conjunction with a contig encoding a mitochondrial ATP/ADP carrier protein, also found in the module. One V-ATPase subunit was found in this module along with clathrin, two proteins that play an important role in diatom cell division and cell wall silicification (45, 46). However, most of the V-type ATPase subunits in the analysis were found in module 2, perhaps implying dynamic regulation of V-type ATPase subunits for different cellular functions throughout different phases of the bloom.

Modules 2 and 5 include transcripts with higher relative abundance at the end of the bloom season, including some transcripts that may be linked to general physiological stress or cell death. Subunits of the proteasome, required for protein degradation and turnover, are enriched in these two modules. Out of 13 proteasome 20S/26S subunits in the WGCNA analysis, eight were found in module 2 or 5 (Fig. S10B). Increased protein turnover versus new protein translation has been described previously in diatom systems as a side effect of decreased growth, cell division and proliferation, and may be associated with nutrient and temperature stress (47). In general, transcripts associated with protein turnover are enriched in these two modules, with 26 of 46 transcripts of relevant KOG class annotation (“Posttranslational modification, Protein turnover, Chaperones”) found in modules 2 and 5 (Fig. S10B). Additional transcripts from modules 2 and 5 associated with protein turnover and related stress response include ubiquitin ligases, aspartyl protease, glutaredoxin and thioredoxin, cyclophilin, and chaperones DnaJ, GroES, and GroEL.

Module 6 includes select transcripts involved in diatom response to nitrogen starvation. Urea transporters and specific ammonia transporters are known to be upregulated in diatoms under nitrogen limiting conditions (48). Both a urea transporter and the *P. australis* homolog of AMT1_1, an ammonia transporter from *P. tricornutum* induced in low-N conditions, can be found in this module. Nitrogen limitation in diatoms is also coupled to an increase in fatty acid biosynthesis and accumulation. Three fatty acid desaturases, including a Δ9-desaturase upregulated in *P. tricornutum* lacking functional nitrate reductase, were also observed in this module (49). Additional genetic markers of diatom physiological state, such as the Si starvation induced silicon transporter *sit1* and the iron-responsive ferritin (FTN) and iron starvation induced proteins (ISIPs), did not meet statistical cutoffs to be included in WGCNA, primarily due to their lack of transcription in a majority of the studied timepoints. This observation is likely due to intermittent induction of these transcripts in direct response to nutrient starvation-induced conditions.

### Episodic summer upwelling perpetuated the lengthy bloom

The decline in *Pseudo-nitzschia* cell counts in early-July (Fig. 1B) coincided with rapid warming and freshening of the mixed layer in Monterey Bay (12). The strong freshening indicated influx of lower-salinity water from the inner California Current, a typical oceanographic response to relaxation of upwelling favorable winds (12, 16). Resurgence of the bloom at MWII in late July likely resulted from resumed upwelling that introduced nutrients to the bay. Following weak upwelling during the first half of July, episodic intensification of upwelling-favorable winds began in late July and persisted for months (12). The resumed influence of upwelling on the bay is evident in satellite SST images. In contrast to warm conditions observed within the bay and at the Point Año Nuevo upwelling center on July 20, cool upwelling plumes originating at Point Año Nuevo were observed to extend into Monterey Bay on July 27 and August 14 (Fig. S11).

### Multiple lines of evidence suggest the *P. australis* bloom was limited by silica and iron

During the relaxation of upwelling in early July, silica concentrations in the water column fell to a historic low and dropped below nitrate concentrations, indicative of silica starvation (12). Silica limitation decreases diatom growth rates and, under starvation conditions, can arrest the cell cycle (50, 51). Diatoms cope with low silica conditions by upregulating silicic acid transporters such as *sit1* to maximize silica uptake (52–54). Indeed, strong induction of *P. australis*-expressed *sit1* was detected in metatranscriptomes in mid-June and early July (Fig. S12), concurrent with the decrease in cell counts from the first phase of the bloom (Fig. 1B). Therefore, elevated *sit1* transcription may be informative for predicting bloom demise under silica-limiting conditions.

Silica limitation can be driven by iron limitation (55). Iron limitation causes diatoms to reduce iron-dependent nitrate uptake and meanwhile continue to take up silica, an iron independent process, thereby drawing down dissolved silica concentrations. Previously unconsidered as a driver of the 2015 bloom, iron limitation was present in the Bay, as inferred by both macronutrient ratios and gene expression. Calculations of Si_ex_, a proxy for diatom iron limitation, suggest pervasive iron limitation throughout the photic zone of Monterey Bay early in the year and alleviation of iron limitation later in the year, as measured in outer Monterey Bay at Station “M1” and inner Monterey Bay at Station “C1” (Fig. 3A, Fig. S13E) (56). Negative Si_ex_ values indicate that diatoms preferentially take up H_4_SiO_4_ relative to NO_3_^-^ due to iron deficiency, and more negative Si_ex_ values indicate a higher level of iron deficiency (56, 57). During the bloom, the mean flow of water in the Bay was from Northwest to Southeast, indicating prevalent advection from M1 toward MWII, and drifter data showed that Monterey Bay was strongly retentive of resident water and phytoplankton biomass (12). M1, in the path of Pt. Año Nuevo’s upwelling plume, is representative of changing conditions that affect the entire bay, including MWII (16). Gene transcription patterns corroborated iron limitation at MWII: genes encoding iron starvation induced protein (ISIP2A) were detected at MWII, indicating iron limitation. *ISIP* expression increased in early-June during the relaxation of upwelling and persisted through the fall (Fig. S14).

**Figure 3.**
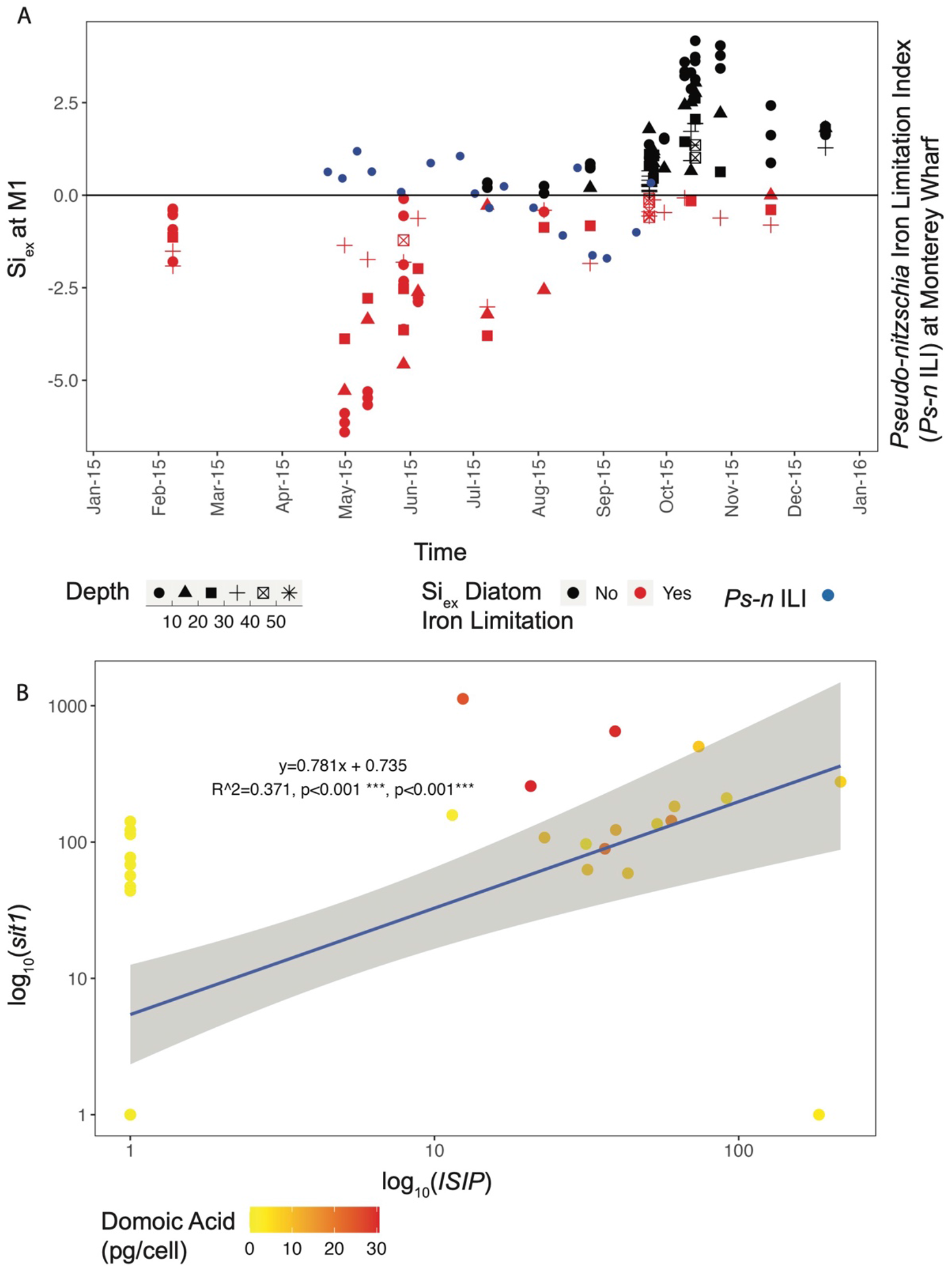
Iron and Si co-limited the bloom and induced toxicity. (A) Negative values for Si_ex_, a proxy for diatom iron limitation, are present during the height of the bloom at Station M1, and co-occur with positive values of the *Pseudo-nitzschia* iron limitation index (*Ps-n* ILI) at Monterey Wharf, indicating iron limitation. As Si_ex_ trends positive later in the year, *Ps-n* ILI trends negative, indicating an alleviation of iron limitation. Si_ex_ was calculated in samples taken within the photic zone above 56 meters depth. (B) *sit1* and *ISIP* are co-expressed and co-occur with higher per cell domoic acid. *ISIP* expression is needed in addition to *sit1* expression for cDA presence. Raw read counts of *sit1* and *ISIP2A* and *ISIP2B* from *Pseudo-nitzschia australis* were normalized to total *Pseudo-nitzschia* read counts, multiplied by 1.0 x 10^6^. A constant of 1 was added to normalized *sit1* and *ISIP*, and the axes were log_10_ transformed.

The resumed upwelling in late-July to early-August was also reflected in the nutrient composition at M1: Si_ex_ began to trend positive indicating an alleviation in iron limitation (Fig. 3A), likely due to a pulse of iron input in the upwelled water. Accordingly, *Pseudo-nitzschia* ferritin (FTN) expression, indicative of long-term iron storage, appeared in August (Fig. S14). *Pseudo-nitzschia’s* FTN usage takes advantage of infrequent pulses of iron, which is beneficial during iron-limited conditions since they can utilize stored iron (58). Furthermore, the *Pseudo-nitzschia* iron limitation index (*Ps-n* ILI), trended from positive to negative in late-July, indicating a switch from iron limited to iron replete *Pseudo-nitzschia* (59) (Fig. 3A). *Ps-n* ILI denotes *Pseudo-nitzschia* iron stress and limitation when the ratio of expression of ISIP2a to FTN is greater than the mean plus one standard deviation of this ratio in iron-replete *Pseudo-nitzschia granii* laboratory cultures (59). Diatom nitrate transporters and ammonium transporters provide further evidence for regional upwelling conditions at MWII: transporter expression peaked during upwelling conditions, both before and after the mid-summer decline in upwelling (Fig. S15).

### Iron and dissolved silica co-limitation actuated the severe bloom toxicity

Our metatranscriptomic data reveal that genes indicative of Si limitation (*sit1*) and iron limitation (*ISIP*) co-occur with each other and with higher per cell DA, suggesting that the *Pseudo-nitzschia* populations experienced nutrient co-limitation, inducing DA production (Fig. 3B). Furthermore, in this dataset, *ISIP* expression must accompany *sit1* expression for cDA quotas to increase; in other words, if *sit1* is highly expressed, but there is no *ISIP* expressed, no DA is measured at the cellular level (Fig. 3B). These data imply that both Si limitation and iron limitation together were critical in inducing the high DA cell quotas observed during the 2015 HAB in Monterey Bay. As Fe’ bioavailability declines, Si:N also declines because silicification of diatoms increases and therefore, silica is limited secondarily (20). This phenomenon of increasing iron limitation and decreasing Si:N ratio has been previously described for diatom populations in the CCE and at a global scale (20, 55, 56). Previously overlooked, iron limitation may have played a significant role in driving the toxicity of the 2015 bloom in Monterey Bay and may have contributed to the Si limitation, both of which likely co-limited the *Pseudo-nitzschia* population.

### Gene transcription predicts domoic acid one week in advance of toxicity

Predicting acute DA events is critical for management of fisheries and for protecting public health. This multi-omic dataset was used as a model for molecular markers to predict DA toxicity. Gene expression is a relevant method for prediction given that transcription occurs temporally prior to protein translation and metabolite production. We found that a generalized linear model of *P. australis* transcripts of *dabA* and *sit1* normalized to total *Pseudo-nitzschia* genus read counts predicts DA production one week in advance of its production (Fig. 4, Fig. S16). Expression of these genes predicts future DA levels a week in advance better than they predict contemporaneous DA levels (Table S5).

**Figure 4.**
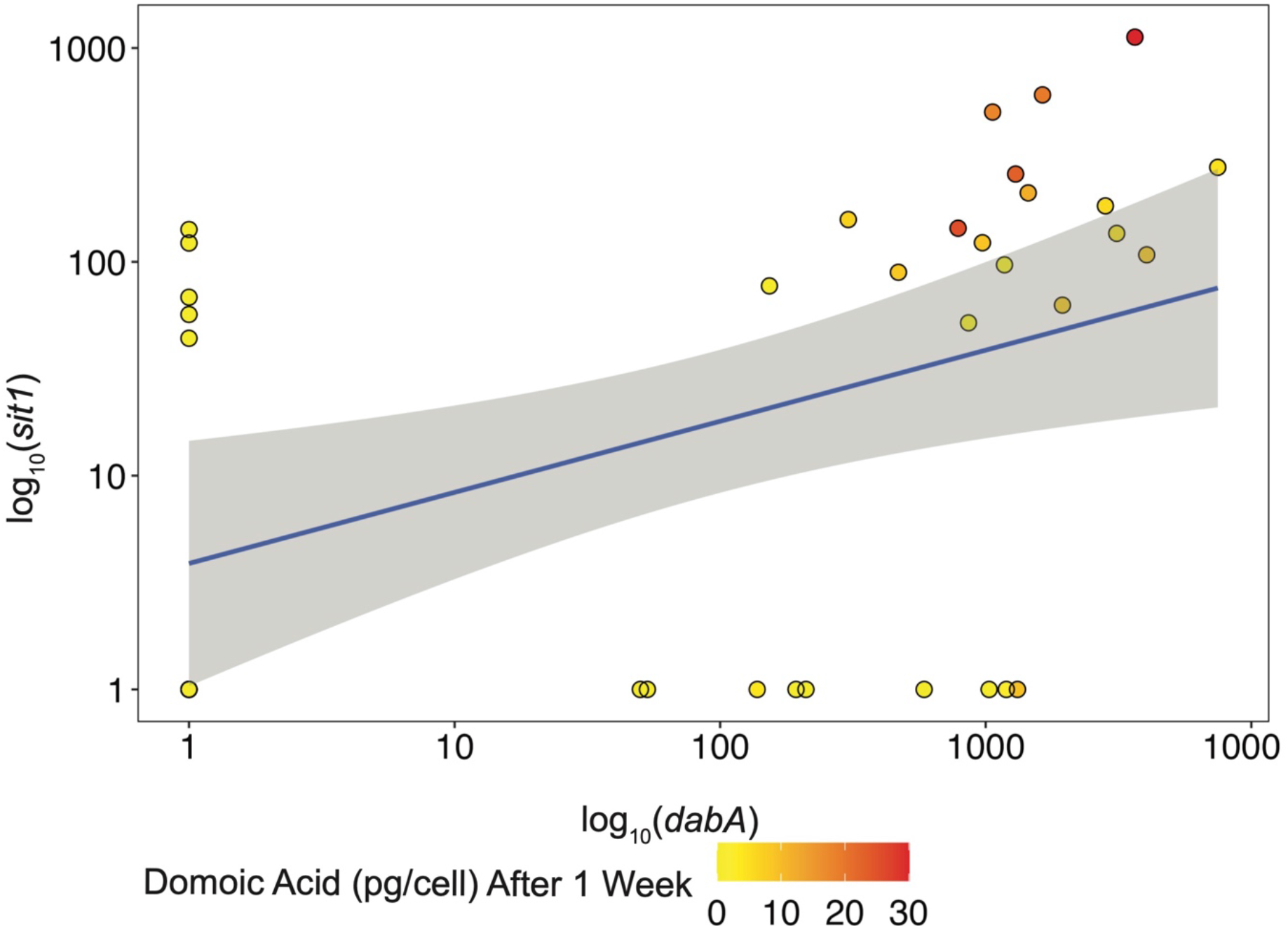
Expression of *sit1* and *dabA* genes predict cellular domoic acid one week in advance of its production. Raw read counts of *sit1* and *dabA* from *Pseudo-nitzschia australis* were normalized to total *Pseudo-nitzschia* read counts, multiplied by 1.0 x 10^6^. A constant of 1 was added to normalized *sit1* and *dabA*, and the axes were log_10_ transformed. Akaike information criterion (AIC) = 236.69, multiple R^2^ = 0.609, adjusted R^2^ = 0.586, p-value = 7.36e-8.

## Conclusions and Discussion

This study provides a robust molecular framework for understanding the progression, toxicity, and physiology of the 2015 *P. australis* HAB event in Monterey Bay, CA. Using 18S, 16S, and ITS2 amplicon sequencing, the transition from a *Chaetoceros*-dominated phytoplankton community to a nearly monospecific bloom of DA-producing *P. australis* was characterized. Due to the iron limiting conditions of the Bay early in the year as inferred by Si_ex_ and *Ps-n* ILI, molecular adaptations by *Pseudo-nitzschia* populations for long-term iron storage under low iron conditions likely enabled *Pseudo-nitzschia* to supersede and outcompete the upwelling-favored diatom *Chaetoceros* (58). Further study will be required to fully understand the competitive advantages that allowed for the dominance of *P. australis* over other toxic and non-toxic *Pseudo-nitzschia* species present before the monospecific bloom, such as *P. multiseries* and *P. fraudulenta*.

Iron limitation most likely contributed to both the competitive success of *Pseudo-nitzschia* and the elevated DA levels observed over several months in the region. Despite conflicting results regarding how iron limitation affects DA production (22, 60–64), we understand that actively growing blooms still in their exponential phase exhibit higher DA production under iron limitation (40). Iron limitation can mediate DA production and release. In other words, iron-stressed *Pseudo-nitzschia* cells are stimulated to produce 10 to 20 times more DA than non-iron-stressed *Pseudo-nitzschia*, and the presence of DA subsequently increases iron uptake by *Pseudo-nitzschia* as demonstrated in both lab and field studies (22, 62). In addition, iron-stressed *Pseudo-nitzschia* have been shown to release 95% of their DA extracellularly (62), a process that has been attributed to the potential role of DA as an iron chelator to bind iron and increase its bioavailability (61). One important caveat is that the relatively low iron-binding constant for DA necessitates an uncommonly elevated concentration of 100 nM of dissolved DA for the molecule to compete with existing iron-binding ligands. Indeed, measurements of dissolved DA around Monterey Bay exceeded 100 nM in our 2015 study (Table S4), strongly suggesting that DA served as a relevant iron-binding ligand and provided *Pseudo-nitzschia* an advantage in acquiring bioavailable iron.

DA events have been increasing in magnitude and intensity over the past several decades for reasons yet to be characterized (2, 65). We hypothesize that the increasing pervasiveness of iron limitation in the CCE is driving both the escalating prevalence of *Pseudo-nitzschia* and the exacerbation of toxin production (56). Global change is altering iron availability and distributions (66). Regions that experience more iron limitation will favor diatoms that use ferritin for long-term iron storage like *Pseudo-nitzschia*, and likely experience heightened DA production (58). Additional lower frequency variability in the CCE may play a role in increasing DA production potential. For instance, in 2015, both the Pacific Decadal Oscillation (PDO) and El Niño Southern Oscillation (ENSO) indices were in their positive phase, resulting in reduced alongshore winds, reduced upwelling, and lower available iron in the euphotic zone, causing the historically low Si_ex_ signature (Fig. 5A, 5B). The high surface and subsurface heat content caused by the unprecedented marine heatwave beginning in 2014 further reduced the strength of upwelling (67, 68) (Fig. 5C). Monterey Bay has also been experiencing an increase in surface pCO_2_ and a decline in pH over the past twenty years, with eventual undersaturation of the system with respect to the carbonate ion and further acidification of upwelled waters due to equilibration with a higher pCO_2_ in the atmosphere as the expected outcome (69). Over the longer term, decreased pH will lower Fe’ uptake rates by ISIP2A, the predominant diatom iron-uptake mechanism, by reducing the availability of the carbonate binding cofactor (70). This trend towards increasing acidification is predicted to intensify iron limitation, and since DA production is correlated with iron limitation and acidification, future world oceans may bear witness to heightened DA (62, 71).

**Figure 5.**
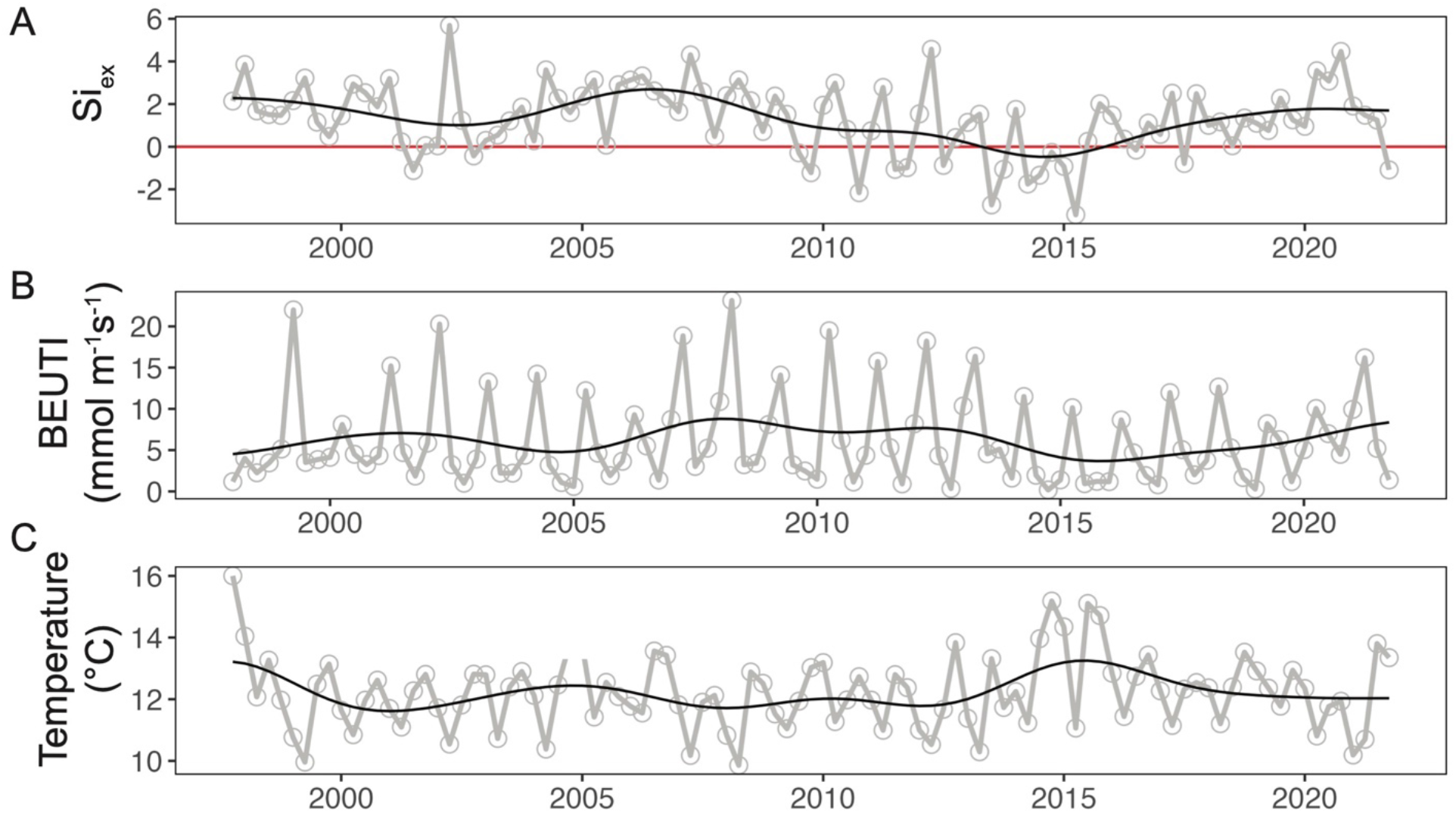
Historical seasonal averages from November 15, 1997 to November 15, 2021 of (top) Si_ex_ at M1 averaged from 0-40 meters, (middle) Biologically Effective Upwelling Transport Index (BEUTI) (https://oceanview.pfeg.noaa.gov/erddap/) from the North American West Coast at 37 °N latitude, (bottom) and temperature (°C) at M1 averaged from 0-40 meters. To calculate Si_ex_, an R preformed value of 0.99 was utilized to estimate an average Si:N of upwelled water in the region. Si_ex_ values below the red line (y=0) indicate time periods of diatom iron limitation.

Active *dab* transcription is inextricably linked to accumulation of DA, and we found that transcription of *<u>dabA</u>* and *sit1* predict DA a week in advance of its production. For optimization of field applications, *P. australis dabA* normalized to the cell counts of *Pseudo-nitzschia “*seriata” size class is a highly predictive and easy-to-use measurement for anticipating DA events with one week lead-time. This unprecedented assay could be effectively deployed by coastal monitoring groups and resource managers as an early warning system that only requires a qPCR assay for *dabA* in addition to the already-collected *Pseudo-nitzschia* cell counts. Nothing like this currently exists, and in fact, many monitoring programs are generally hindered by long lags in chemical analysis of DA that limit the benefits of real-time observing. Metatranscriptomic sequencing of *Pseudo-nitzschia* bloom events, including more frequent sampling on the timescale of days rather than weeks, will be necessary to fine-tune the timeline for *dab* detection to better inform bloom forecasting. Additionally, the precise nucleotide sequence of *dab* transcripts lends insight into DA producing species. Given the paucity of *dab* transcripts currently identified, further sequencing of *dab* genes and transcripts will be required to fully explore this trend among all DA-producing *Pseudo-nitzschia* species and isolates. Additionally, induced expression of silica-responsive genes such as *sit1* may reflect active diatom silica limitation or starvation and may help predict impending bloom demise.

The metatranscriptomic approach employed in this study has also enabled preliminary insight into the changing physiology of *Pseudo-nitzschia* HABs from inception to demise. We described trends in the relative expression of genes involved in active growth, photosynthesis, toxin production, and environmental stress. A targeted, Lagrangian sampling of surface waters or subsurface chlorophyll maxima throughout a bloom event may yield improved functional clustering of gene expression by following discrete *Pseudo-nitzschia* subpopulations in the water column (72). Models for *Pseudo-nitzschia* HAB physiology will be improved further if Lagrangian sampling is conducted with higher frequency.

This study represents the most comprehensive molecular description of any eukaryotic marine HAB event. This work connects routine monitoring data, such as toxin measurements and phytoplankton cell counts to molecular HAB species identification and toxin biosynthesis gene expression. The synthesis of these data provides new insights into *Pseudo-nitzschia* HAB physiology as it relates to nutritional and physical conditions and provides an improved molecular framework for HAB monitoring and prediction. The continued study of molecular physiology in HABs promises to lend essential insight into the nature of toxin production and bloom formation in the changing world ocean.

## Materials and Methods

Method details are available in SI Appendix. All routine monitoring data collected weekly from Monterey Municipal Wharf II (MWII) during the study period, including *Pseudo-nitzschia* cell counts, DA measurements, local chlorophyll concentration, temperature, and nutrient data, is made publicly available through the Southern California Coastal Ocean Observing System (SCCOOS, https://sccoos.org/harmful-algal-bloom/) as part of the California Harmful Algal Bloom Monitoring and Alert Program (CalHABMAP).

RNA was extracted using a modified version of the Direct-zol RNA Miniprep Plus kit (Zymo Research), and cDNA was synthesized using the SuperScript III First-Strand Synthesis System (Life Technologies). Poly-A enriched RNA was synthesized using the TruSeq Stranded mRNA prep kit (Illumina). Amplicons were sequenced on the MiSeq PE300 (Illumina), and the poly-A enriched RNA on the HiSeq4000 PE150 platform (Illumina). Sequence data have been deposited in the NCBI Sequence Read Archive under accession number PRJNA1027375.

## Acknowledgements

We thank the Southern California Coastal Ocean Observing System (SCCOOS) and California Harmful Algal Bloom Monitoring and Alert Program (HABMAP) for data collection and availability (NOAA Award No. NA21NOS0120088 and NA11NOS0120032). We thank Jeffrey B. McQuaid and Zoltan Fussy (both J. Craig Venter Institute) for helpful discussions. This research was supported by the National Oceanic and Atmospheric Administration (NA19NOS4780181 to A.E.A., B.S.M., J.P.R., and C.R.A., and NA11NOS4780030 and to R.M.K.), the National Institutes of Health (F31ES030613 to J.K.B.).

## Supplementary Information

## Materials and Methods

### Routine Monitoring Data Availability and Visualization of Satellite Data

All routine monitoring data collected weekly from Monterey Municipal Wharf II (MWII) during the study period, including *Pseudo-nitzschia* cell counts, DA measurements, local chlorophyll concentration, temperature, and nutrient data, are made publicly available through the Southern California Coastal Ocean Observing System website (SCCOOS, https://sccoos.org/harmful-algal-bloom/) as part of the California Harmful Algal Bloom Monitoring and Alert Program (CalHABMAP). Satellite data were acquired and visualized via the Environmental Research Division’s Data Access Program (ERDDAP) data server from the National Oceanic and Atmospheric Association (NOAA). All-surface temperature data were collected in Local Area Coverage (LAC) format by the Advanced Very High Resolution Radiometer (AVHRR) scanner onboard NOAA’s Polar Orbiting Environmental Satellite (POES). Satellite-measured remote chlorophyll concentration (mg/m^3^) was collected by the Visible Infrared Imaging Radiometer Suite (VIIRS) onboard the Suomi National Polar-Orbiting Partnership (Suomi NPP) satellite.

### Phytoplankton Net Tow and Sample Filtering

Weekly net tows from MWII were obtained using a 20 cm diameter, 20 μm mesh net to concentrate surface waters to a depth of five meters, as is routine for monitoring at MWII. From the net tows, 100 mL of concentrated phytoplankton samples were applied to 0.22 μm polycarbonate (PC) filters, which were then stored in 1 mL of TRIzol (Invitrogen) reagent in 15 mL conical tubes, mixed by vortexing, and kept at -80°C for future processing. A total of 53 samples were processed and stored in this manner.

### RNA extraction and cDNA generation

Samples were removed from -80°C storage. The PC filter was removed from the TRIzol solution and samples were subsequently centrifuged for 20 minutes at 2500 x g and 4 °C to pellet particulate matter. The supernatant was then transferred to a new tube. RNA extraction begins with the addition of 200 μL of chloroform to each sample, which were then shaken vigorously for 15 seconds and allowed to stand for 12–15 minutes at room temperature (∼25 °C). The resulting mixture was then centrifuged at 12,000 xg for 15 minutes at 4 °C. Next, the aqueous phase was transferred to a new tube, adding an equal volume of 100% ethanol and mixing well. The mixture was then transferred to a Zymo-Spin IIICG Column (Zymo Research), after which the RNA extraction was completed using the Direct-zol RNA Miniprep Plus kit (Zymo Research) following the manufacturer protocol, including an on-column DNA digestion step using DNase I (New England Biolabs). Suitable RNA quality was verified using an Agilent 2100 Bioanalyzer.

Synthesis of cDNA to be used as template for generation of 18S-V4, 16S and ITS2 amplicons was preformed using the SuperScript III First-Strand Synthesis System (Life Technologies). Following kit standard protocol, 100 ng of total RNA per sample was used with random hexamer primers to make 20 μL of cDNA.

### Preparation and sequencing of 18SV4, 16S, and ITS2 amplicon libraries

Amplicon libraries for 18SV4, 16S and ITS2 sequences were generated using the cDNA libraries from above as a template for the one-step PCR reactions to simultaneously amplify target sequences and incorporate Illumina adaptors, linker sequences, and unique barcoded indices. All one-step PCR reactions were performed using the TruFi DNA Polymerase PCR kit (Azura). All primers used in this study, together with attached adaptors, linkers and barcoded indices are listed in Table S1. The 18SV4 region was amplified using the primer pair V4F and V4RB (1). The 16S(V4-V5) region was amplified using the primer pair 515F-Y and 926R (2). The ITS2 region was amplified using an unpublished primer pair 5.8SF and 28SR. Primer pairs containing unique combinations of barcoded indices, allowing downstream demultiplexing, are generated in 10 μM concentration and stored in 96-well plates.

PCR reactions are then set up using 1 μL of cDNA as template, 0.4 μM of primer pair, and TruFi DNA Polymerase and buffer mix per manufacturer protocol and brought to a total reaction volume of 25 μL using molecular grade water. PCR reactions for 18SV4 and 16S amplicon generation were performed with an initial denaturing step at 95°C for one minute followed by 30 cycles of denaturing at 95°C for 15 seconds, annealing at 56°C for 15 seconds, and extension at 72°C for 30 seconds. Initial PCR reactions for ITS2 followed a similar protocol, however improved amplification was observed when the annealing temperature was decreased to 51-53°C. To confirm amplification and correct size of amplicon, 2.5 µL of each PCR reaction was run on a 1.8% agarose gel.

The PCR products were then cleaned up using AMPure XP beads (Agencourt) and the standard PCR purification protocol per manufacturer recommendations and eluted in 35 µl of elution buffer. Cleaned PCR products were then quantified using the PicoGreen Quant-IT dsDNA Quantitation Reagent (Life Technologies). Equal quantities of 18SV4, 16S or ITS2 amplicons (∼10 ng per reaction) were pooled, cleaned, and concentrated using AMPure XP beads using standard protocol, eluting in 45 µl of elution buffer. Final library quality was assessed using a TapeStation (Agilent) and quantified on a Qubit fluorometer (ThermoFisher). Respective pools were then sequenced on the MiSeq PE300 (Illumina) generating 250 bp paired end reads for all amplicon libraries. Sequencing was performed at the UC Davis Sequencing Core.

### Assembly, taxonomic classification and visualization of amplicon sequencing

The 18SV4, 16S and ITS2 amplicon sequences were imported separately into Qiime2 where the dada2 plug-in was used to merge, quality filter the reads, identify and remove chimeric reads, and generate Amplicon Sequence Variants (ASVs) and count tables (3, 4). Taxonomy was assigned to the 18S ASVs using a qiime2 naïve-bayes classifier trained on the PR2 version 4.11.1 database (5). The 16S ASVs were annotated first using a qiime2 naïve-bayes classifier trained on the Silva-132 database in order to differentiate bacterial and mitochondrial sequences from chloroplast sequences (6). Chloroplast ASVs were extracted in Qiime2 and further annotated using a qiime2 naïve-bayes classifier trained on the PhytoREF database (7). Genus and species classifications for ITS2 ASVs were assigned using the “classify-consensus-blast” feature classifier, using a database of *Pseudo-nitzschia* ITS2 sequences as the reference BLAST database (8). Conglomeration of ASV counts by taxonomic level and subsequent generation of stacked bar plots was performed in PhyloSeq v.1.34.0 (9).

### Preparation and sequencing of polyA-enriched RNA

Starting with 500 ng of total RNA per sample, we used the TruSeq Stranded mRNA prep kit (Illumina) to make polyA-enriched libraries suitable for RNA sequencing, following manufacturer protocols. Following preparation of polyA-enriched libraries, samples were combined into two pools and ran on the HiSeq4000 PE150 platform (Illumina) at the UC Davis Sequencing Core.

### Assembly and annotation of RNA sequencing libraries

The resulting demultiplexed HiSeq4000 read libraries were processed via the RNAseq Annotation Pipeline v0.4 (10). Reads were first trimmed for quality and filtered to remove primers, adaptors and rRNA sequences using Ribopicker v.0.4.3 (11). CLC Genomics Workbench 9.5.3 (QIAGEN) was used to assemble contigs, first by library, then globally. Open reading frames (ORFs) were predicted from the assembled contigs using FragGeneScan (12). The resulting ORFs were annotated *de novo* for function and taxonomy via KEGG, KO, KOG, Pfam, TigrFam and the reference dataset PhyloDB 1.02 (10). Reads were mapped back to assembled contigs using BWA MEM to generate read counts (13).

### Defining expression modules for the *P. australis* bloom

The Weighted Gene Correlation Network Analysis (WGCNA) R package was implemented to identify modules of *P. australis* ORFs with similar expression patterns to define functional clusters (14). The bloom period for WGCNA analysis purposes was defined as 15 April through 30 September of 2023. *P. australis* ORFs were identified and filtered based on taxonomic annotation above, leveraging the PhyloDB 1.02 database which contains a *P. australis* annotated transcriptome (10, 15). This yielded 14873 *de novo* assembled *P. australis* ORFs. To obtain statistically relevant results, only ORFs with 10 or more reads in 80% of the libraries comprising the bloom period were considered, yielding 437 ORFs for WGCNA analysis.

Following the filtering step, expression was normalized by dividing reads mapped to a given ORF by the total number of reads mapped to all assembled *P. australis* ORFs to account for library and *P. australis* population size prior to WGCNA analysis and construction of a correlation matrix. An adjacency matrix was built from the correlation matrix input by applying a power function (AF(s)=s^b) to the input data, where “b” is defined as the soft-thresholding parameter. A soft-thresholding parameter of b = 6 was found to be lowest value at which a scale-free topology R^2^ value exceeding 0.8 was achieved (Fig. S17). This information was used to construct the consensus matrix and dendrogram using the function “blockwiseConsensusModules” using the parameters: b = 6, TOMType = “signed”, detectCutHeight = 0.995, reassignThreshold = 0 (Fig. S18). The “moduleEigengenes” function was used with default parameters to select an “eigengene,” or the ORF whose expression profile is most representative of the other profiles contained in the module. No minimum module size was set and no modules were merged to generate modules containing genes with the highest possible similar expression profiles to one another, maximizing the variance explained by each eigengene.

### Analysis of markers for iron limitation and DA prediction

Si_ex_ was calculated as the concentration of silicate minus the concentration of nitrate times the ratio of H_4_SiO_4_ to NO_3_^-^ at the regional upwelling source depth (16). Si_ex_ was calculated assuming that the ratio (μmol/L:μmol/L) of H_4_SiO_4_ to NO_3_^-^ at the regional upwelling source depth is equal to 1.

*Pseudo-nitzschia* iron limitation index (*Ps-n* ILI) was calculated from equations based on the expression of ISIP2a relative to FTN of laboratory-grown *P. granii* cells (17). *Ps-n* ILI calculations were preformed using normalized gene expression in transcripts per million (TPM) rather than raw read counts. Samples were not omitted if ferritin expression was zero.

Linear regressions were conducted using glm() and lm() functions in the stats package (version 3.6.2) in RStudio (version 2022.02.3). Gene expression raw read counts from *Pseudo-nitzschia australis* were normalized to total *Pseudo-nitzschia* read counts for analysis.

## Supplemental Table and Figures

**Figure S1.**
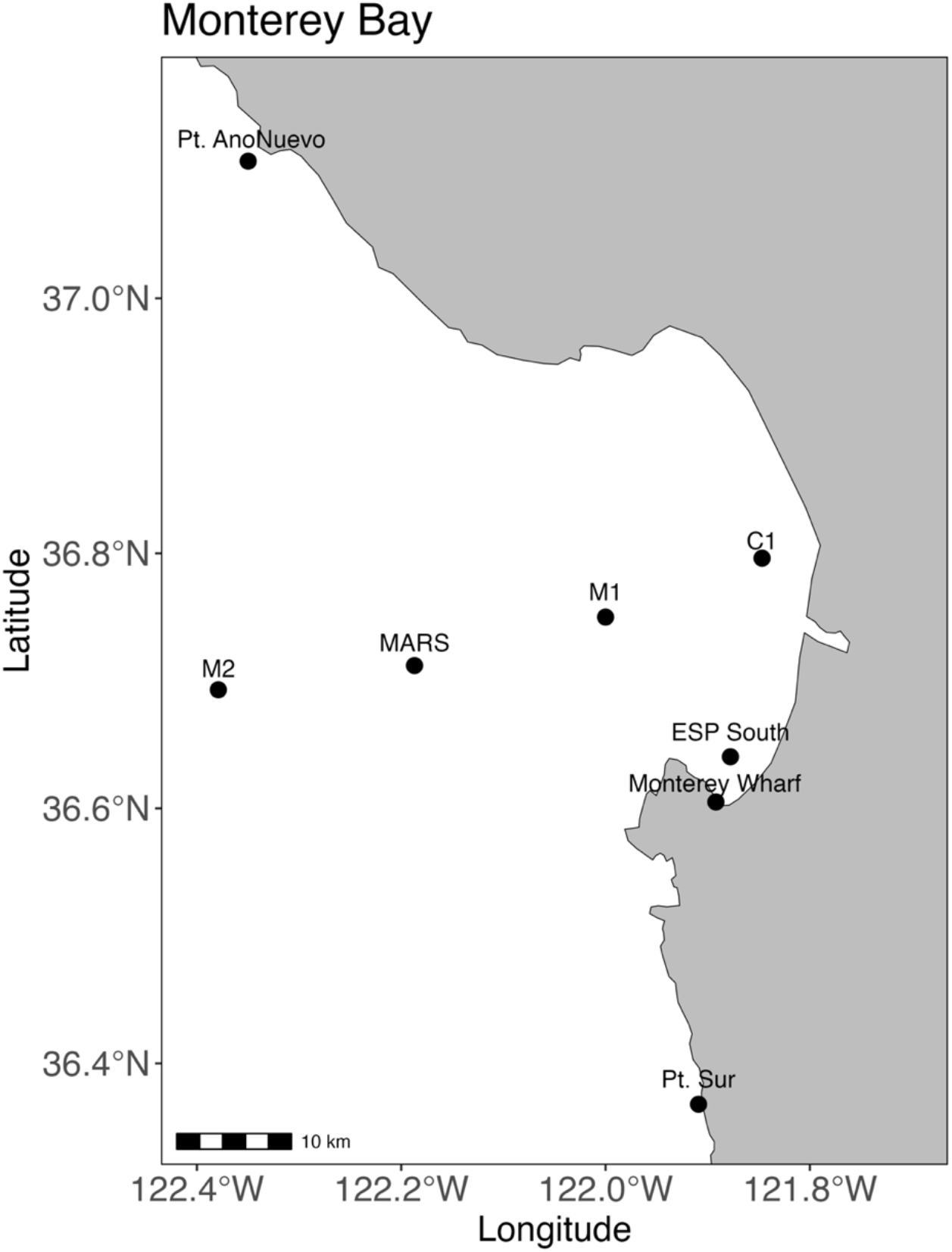
Sampling location at Monterey Wharf II (MWII) is marked together with the upwelling centers outside the bay, Pt. Año Nuevo and Pt. Sur. Also marked are regional monitoring locations used for Si_ex_ analysis.

**Figure S2.**
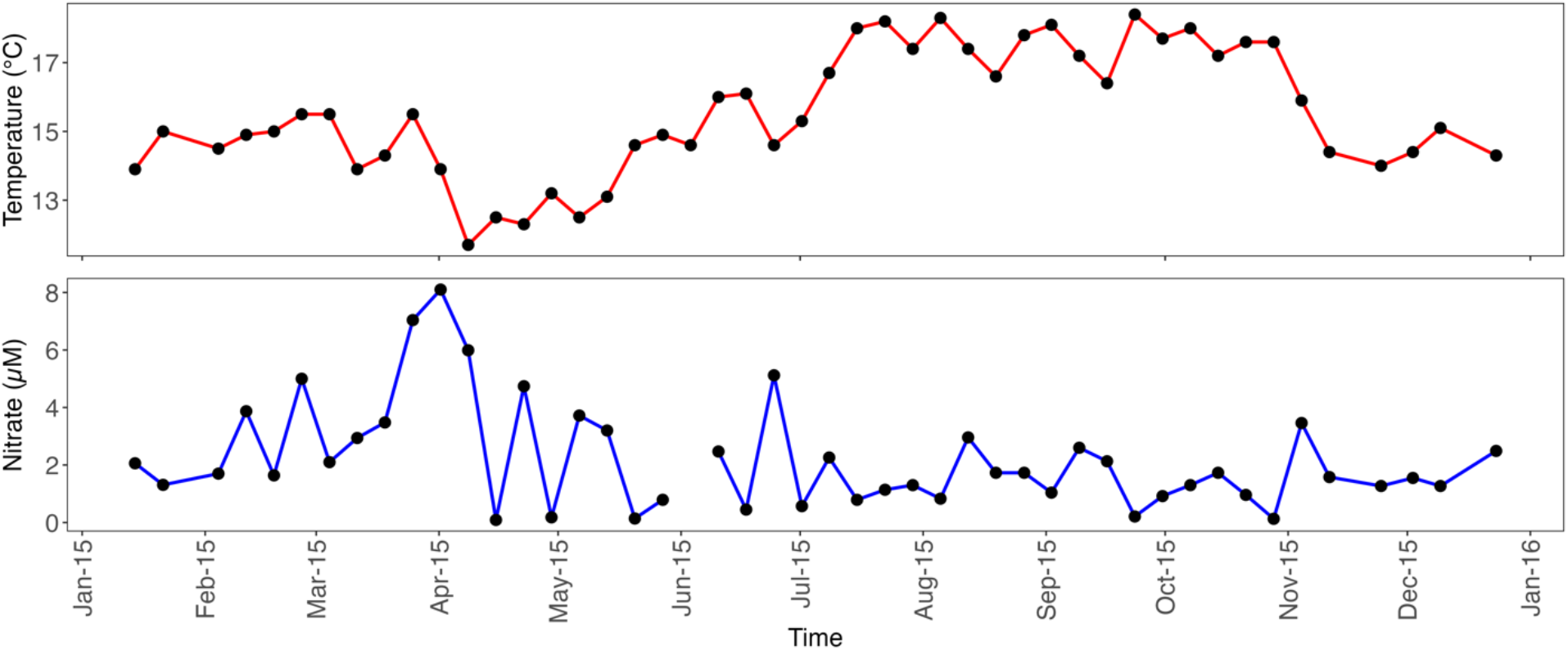
(Top) Temperature (°C) and (Bottom) nitrate (μM) measurements collected from MWII.

**Figure S3.**
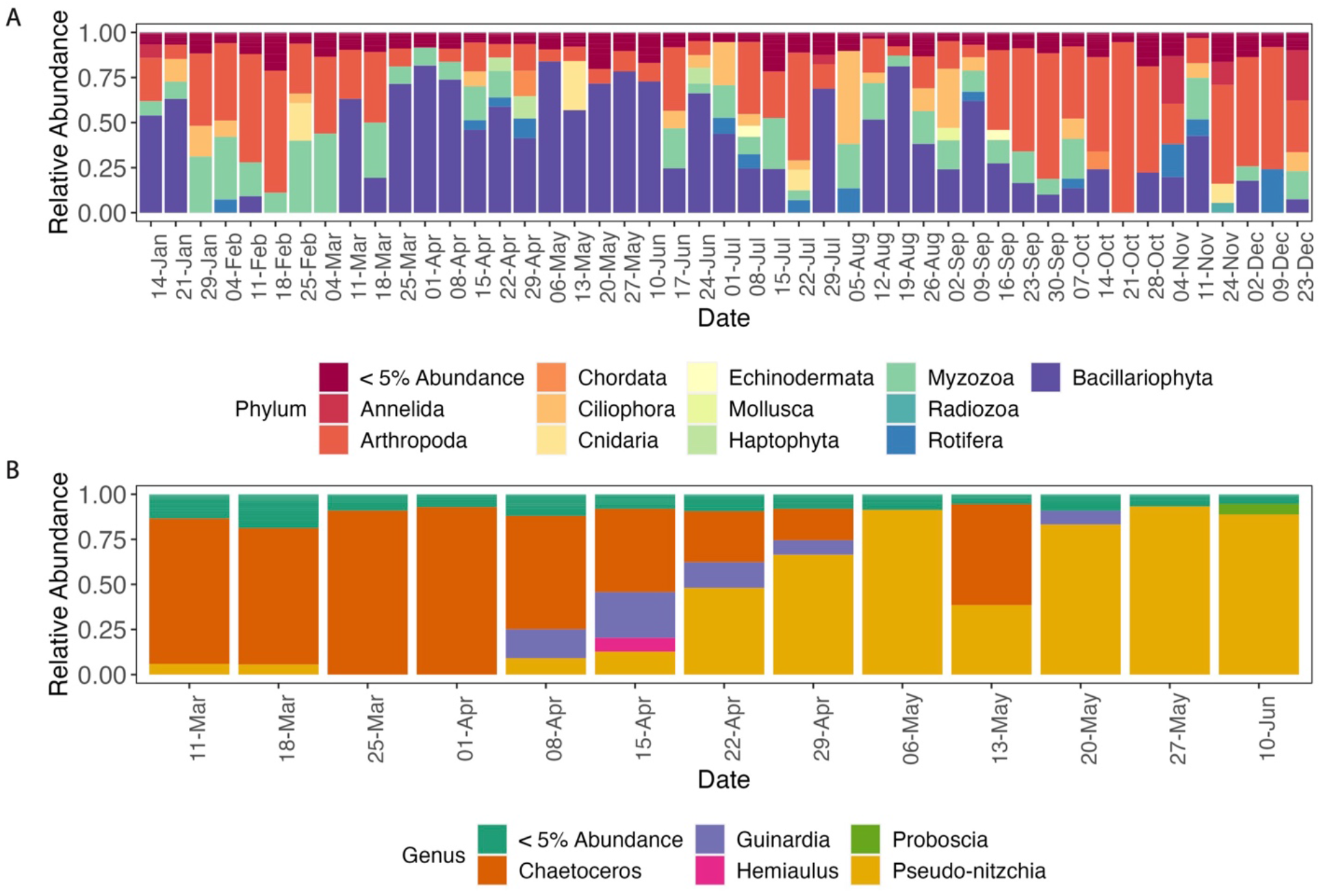
18S-V4 Amplicon Sequencing of 2015 MWII Samples. (A) Relative abundance of phyla in 18S-V4 amplicon sequencing libraries, (B) Relative abundance of diatom (Bacillariophyta) genera during the spring phytoplankton succession.

**Figure S4.**
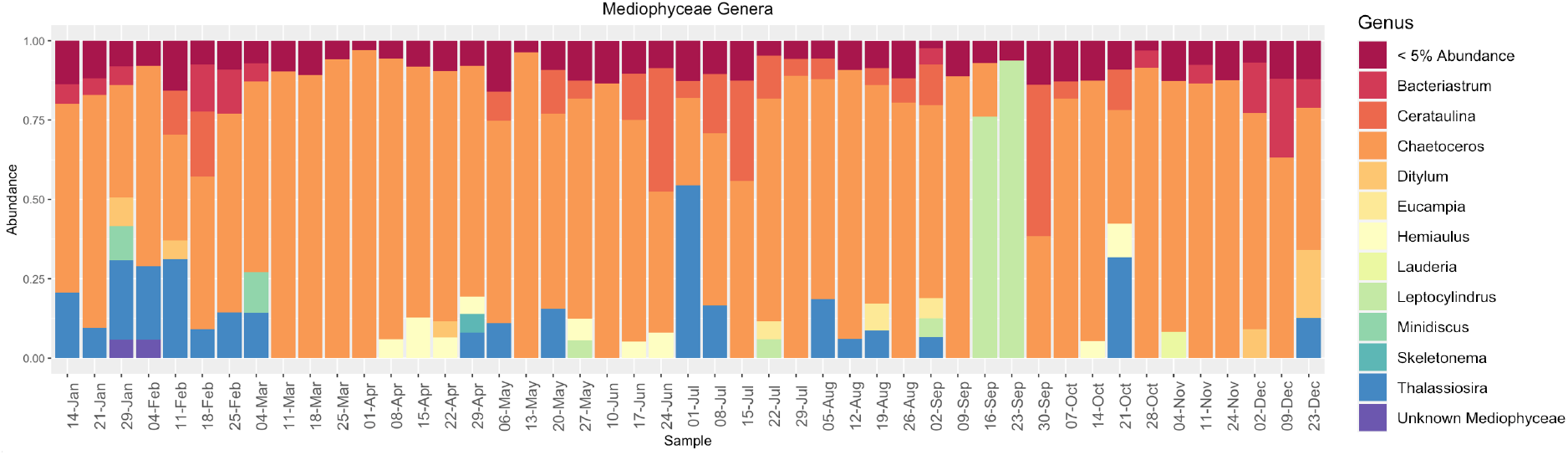
Genus composition of Mediophyceae (centric) diatoms determined by 18SV4 sequencing from weekly samples throughout 2015

**Figure S5.**
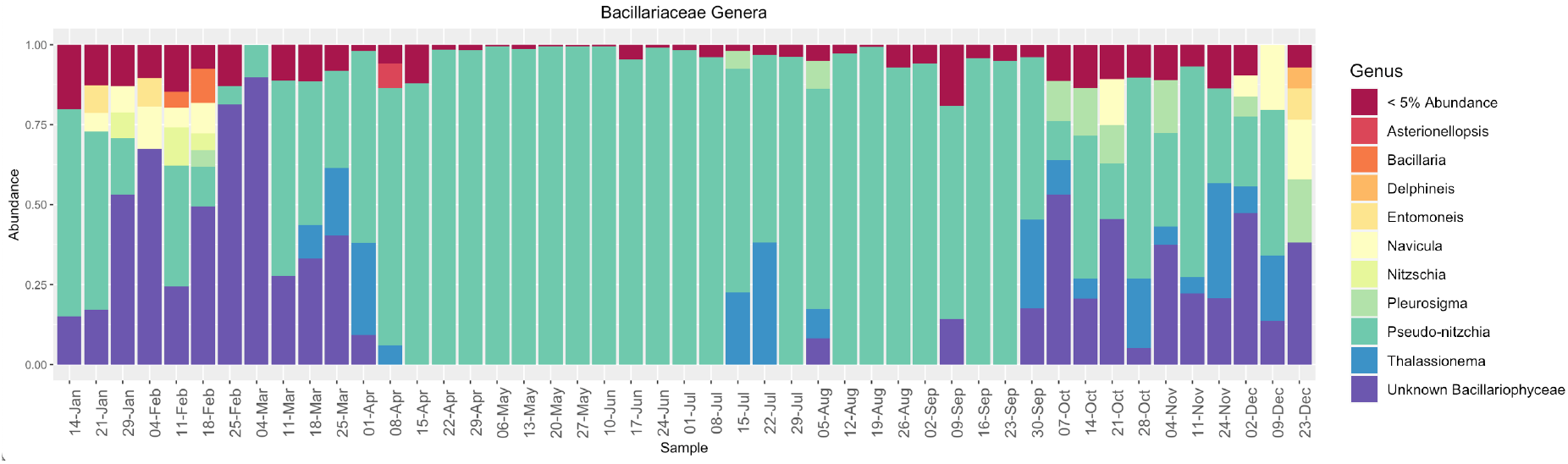
Genus composition of Bacillariophyceae (pennate) diatoms determined by 18SV4 sequencing from weekly samples throughout 2015.

**Figure S6.**
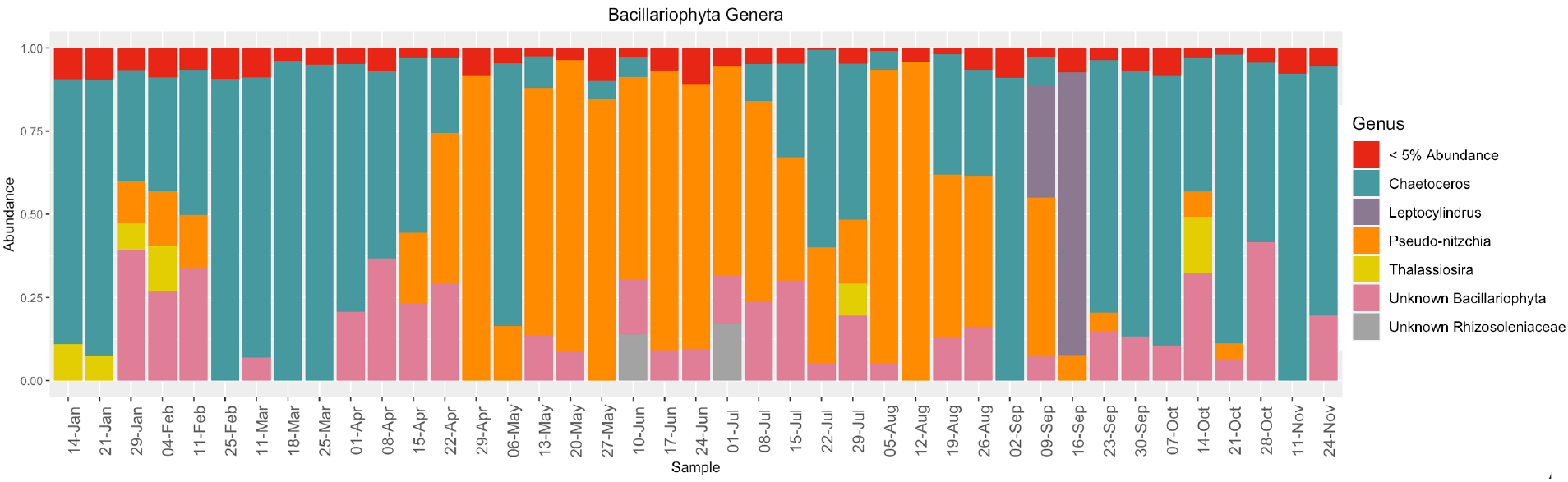
Genus composition of diatoms (Phylum Bacillariophyta) determined by 16S chloroplast sequencing from weekly samples throughout 2015.

**Figure S7.**
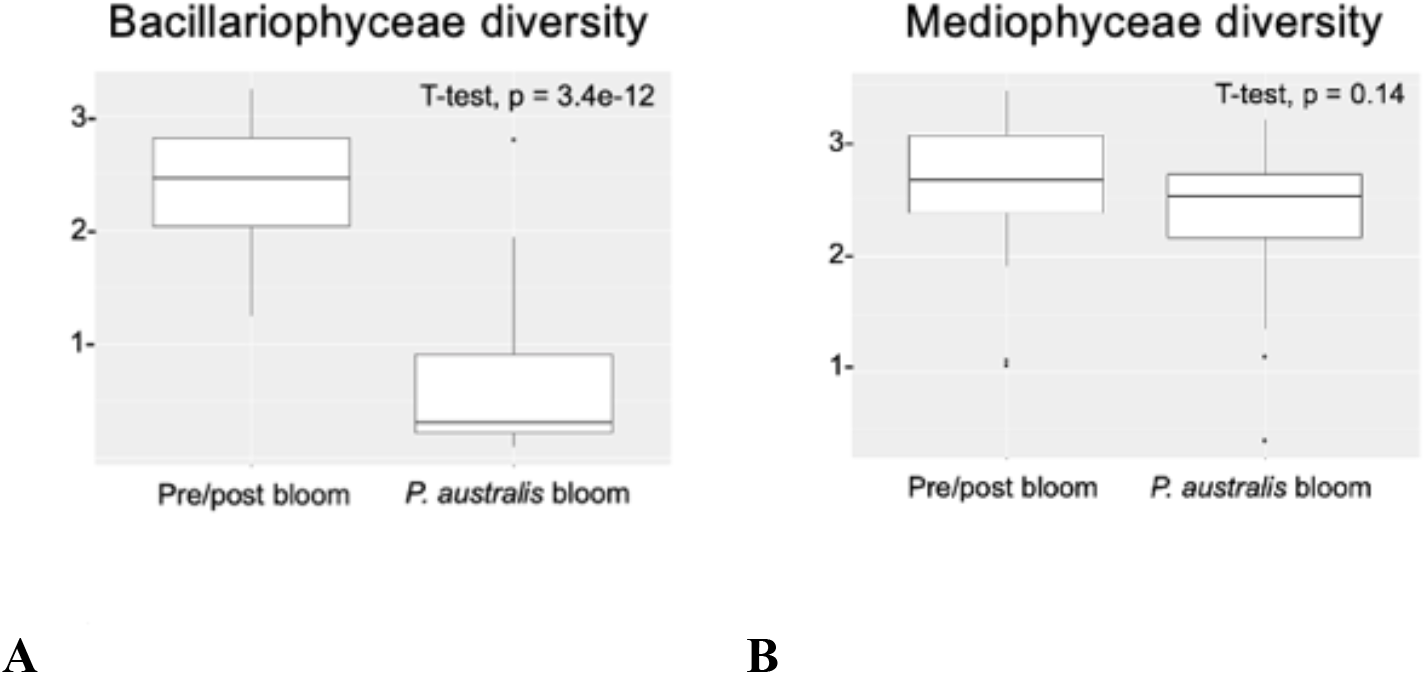
18S-V4 Amplicon Sequencing of 2015 MWII Samples. (A) Shannon alpha-diversity of pennate (Bacillariophyceae) diatoms comparing *P. australis* bloom samples (Apr 22^nd^ – Sep 30^th^) with non-bloom samples (rest of year). (B) Shannon alpha-diversity of centric (Mediophyceae) comparing bloom samples (Apr 22^nd^ – Sep 30^th^) with non-bloom samples (rest of year).

**Figure S8.**
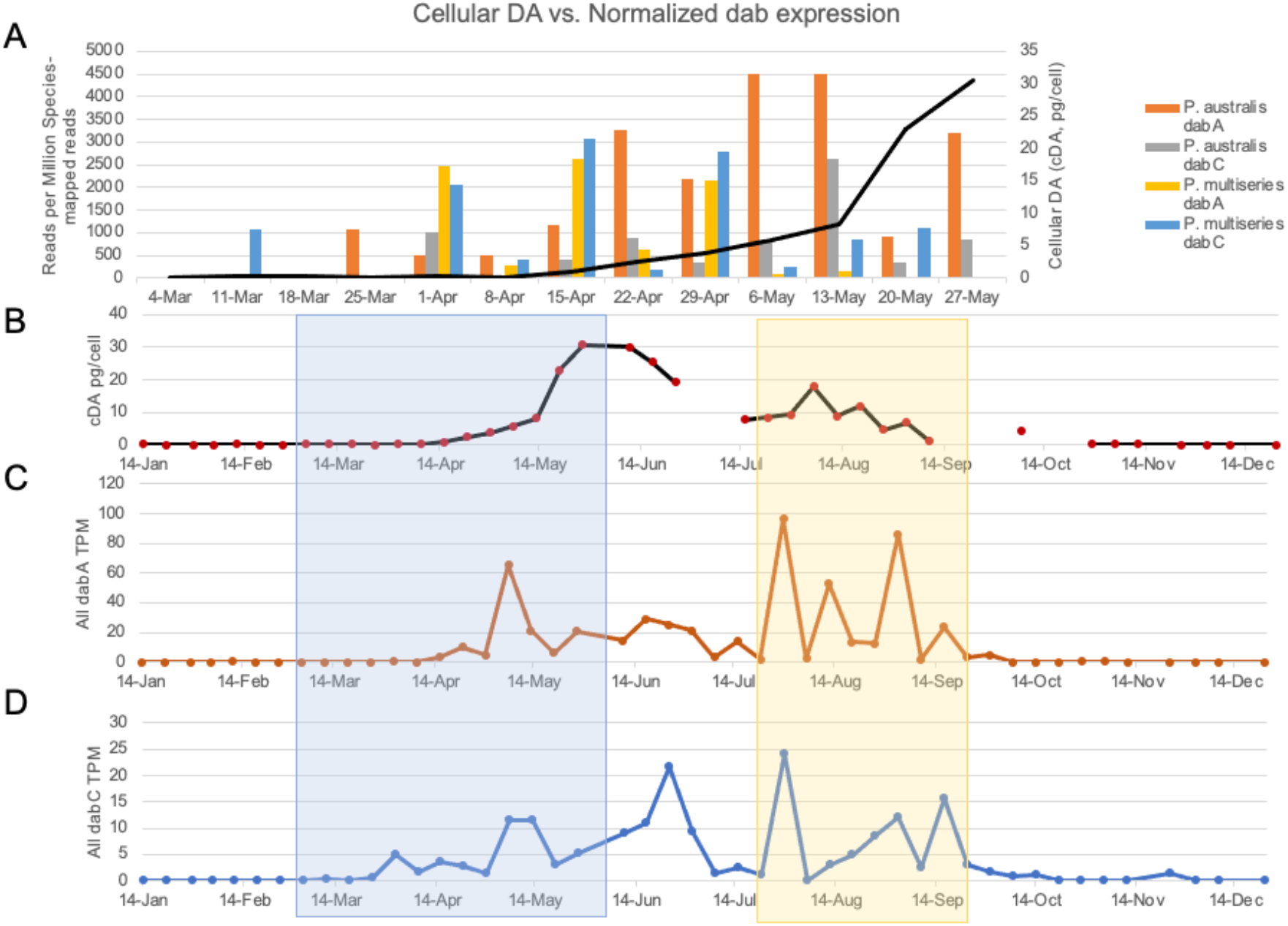
Transcription of *dabA* and *dabC* throughout the bloom. (A) Normalized expression of *dabA* and *dabC* transcripts plotted against cellular DA (cDA, pg/cell) during bloom initiation in the spring. Assigned reads for *P. australis* or *P. multiseries dab* transcripts were normalized by the total number of *P. australis* or *P. multiseries*-mapped reads per library to estimate relative expression within each DA-producing species, (B) cDA values for the 2015 calendar year, (C) Summed expression of all *dabA* transcripts, represented as library normalized per-million reads, (D) Summed expression of all *dabC* transcripts, represented as library normalized per-million reads. Early and late bloom phases indicated in (B-D) with blue and orange boxes, respectively.

**Figure S9.**
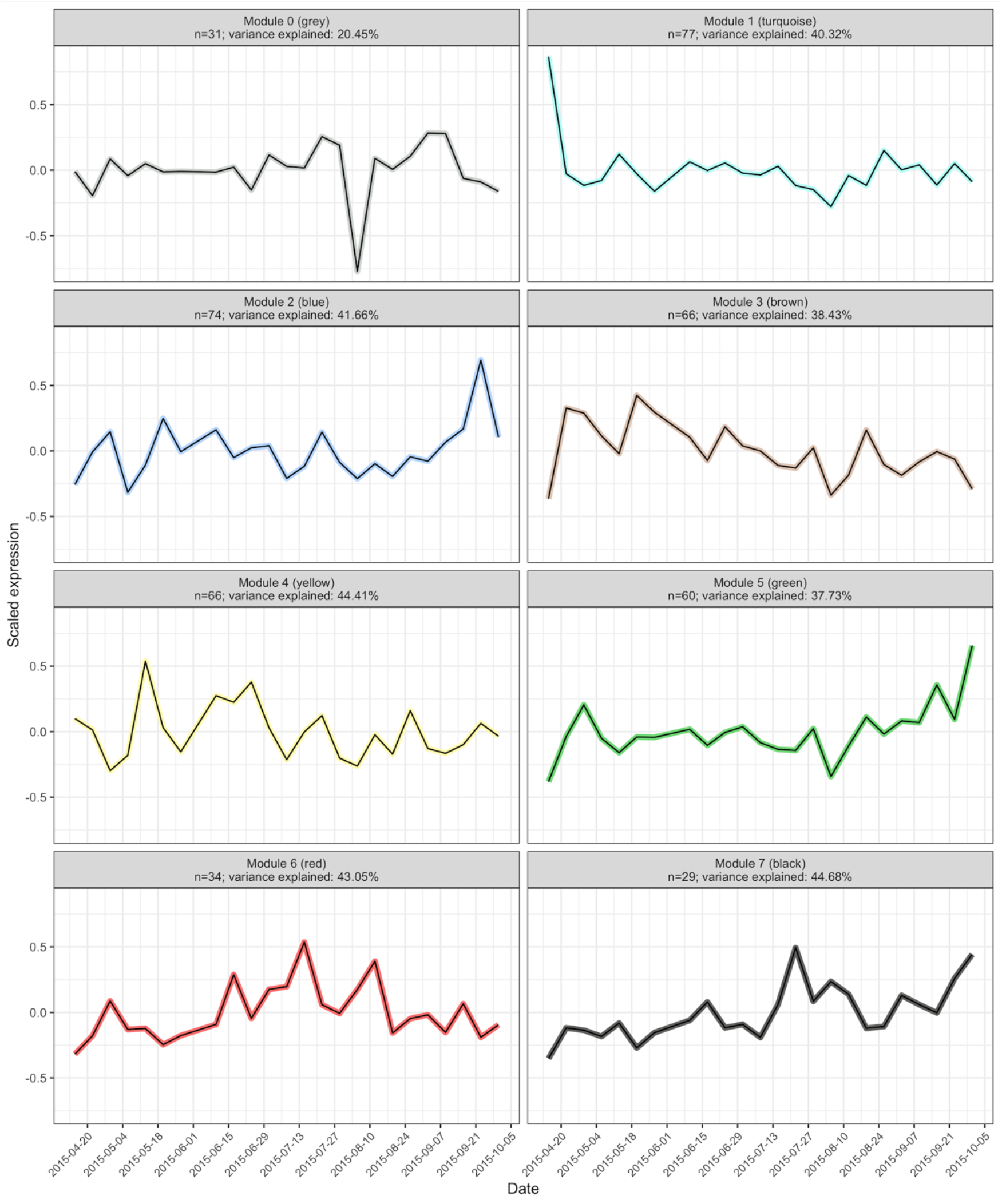
Relative Expression Modules for *P. australis* transcripts throughout the HAB event determined by WGCNA. Weighted gene correlation network analysis (WGCNA) on a highly-expressed subset of the *de-novo* assembled *P. australis* HAB metatranscriptome identified seven modules of transcripts with similar expression profiles (modules 1-7), including one module of remaining contigs with low overall variance explained by the model (module 0).

**Figure S10.**
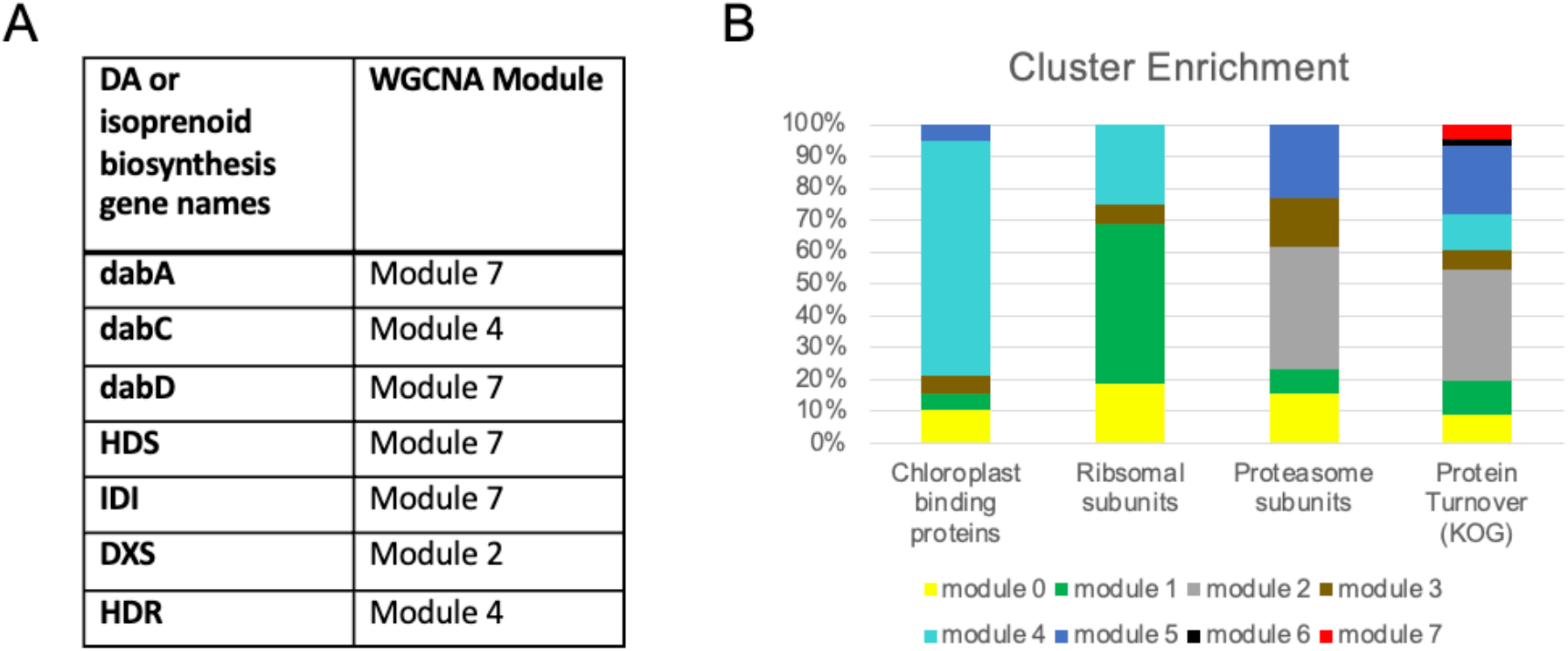
Annotations of interest from WGCNA clustering analysis (*A*) Functional clustering of DA biosynthesis (*dab*) and isoprenoid biosynthesis transcripts. (*B*) Enrichment of ORF annotations in WGCNA modules. Chloroplast binding proteins, ribosomal subunits and proteasome subunits were determined on the basis of PFam annotation, among other annotations. “Protein 8urnover (KOG)” ORFs includes all proteins in the KOG class “Posttranslational modification, Protein turnover, Chaperones.”

**Figure S11.**
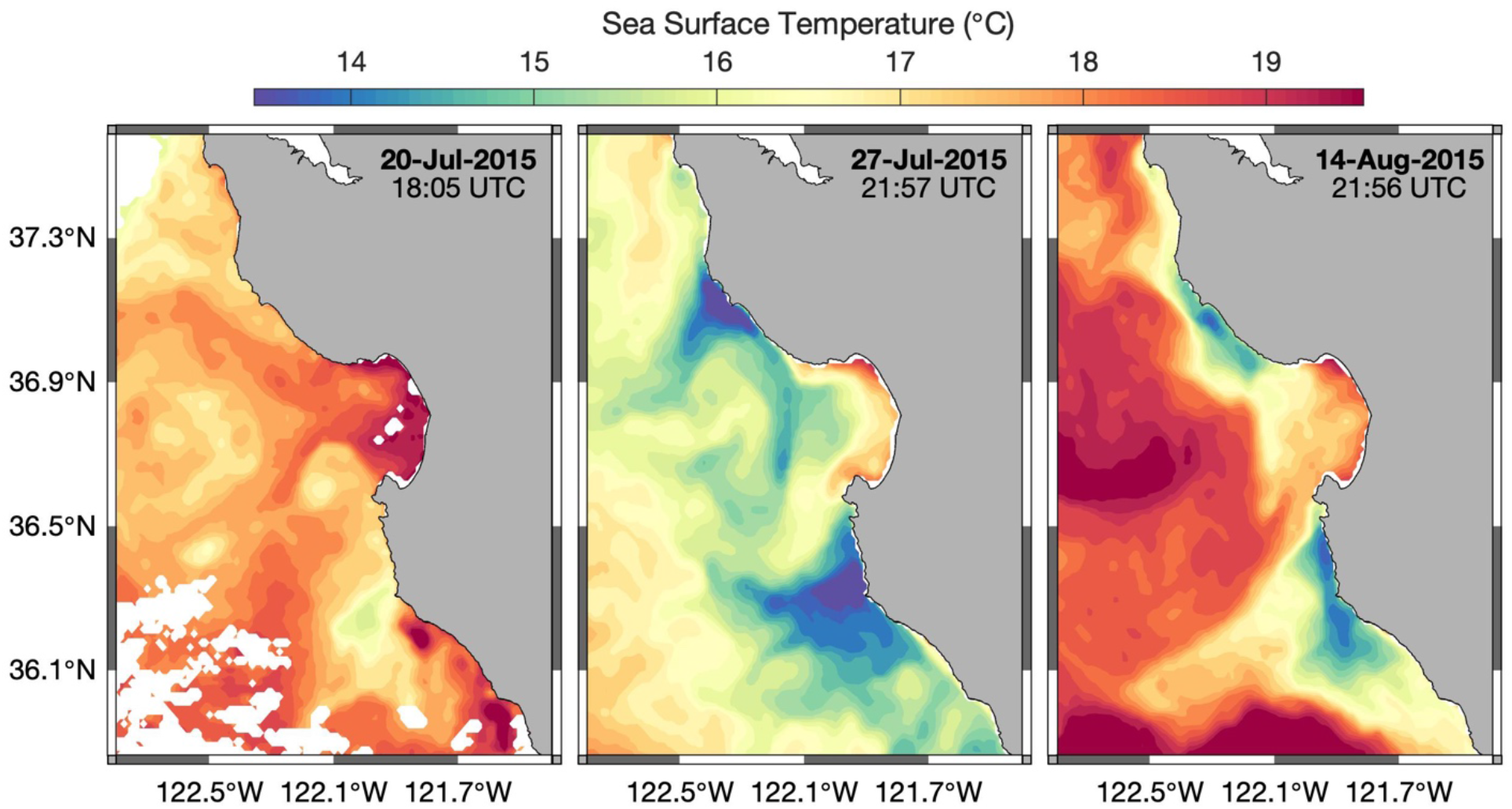
Satellite imagery of Monterey Bay illustrating upwelling (cool plumes) during late-July and mid-August. Sea surface temperature (°C) of Monterey Bay on (left) 20-July (middle) 27-July, and (right)14-August from NOAA AVHRR.

**Figure S12.**
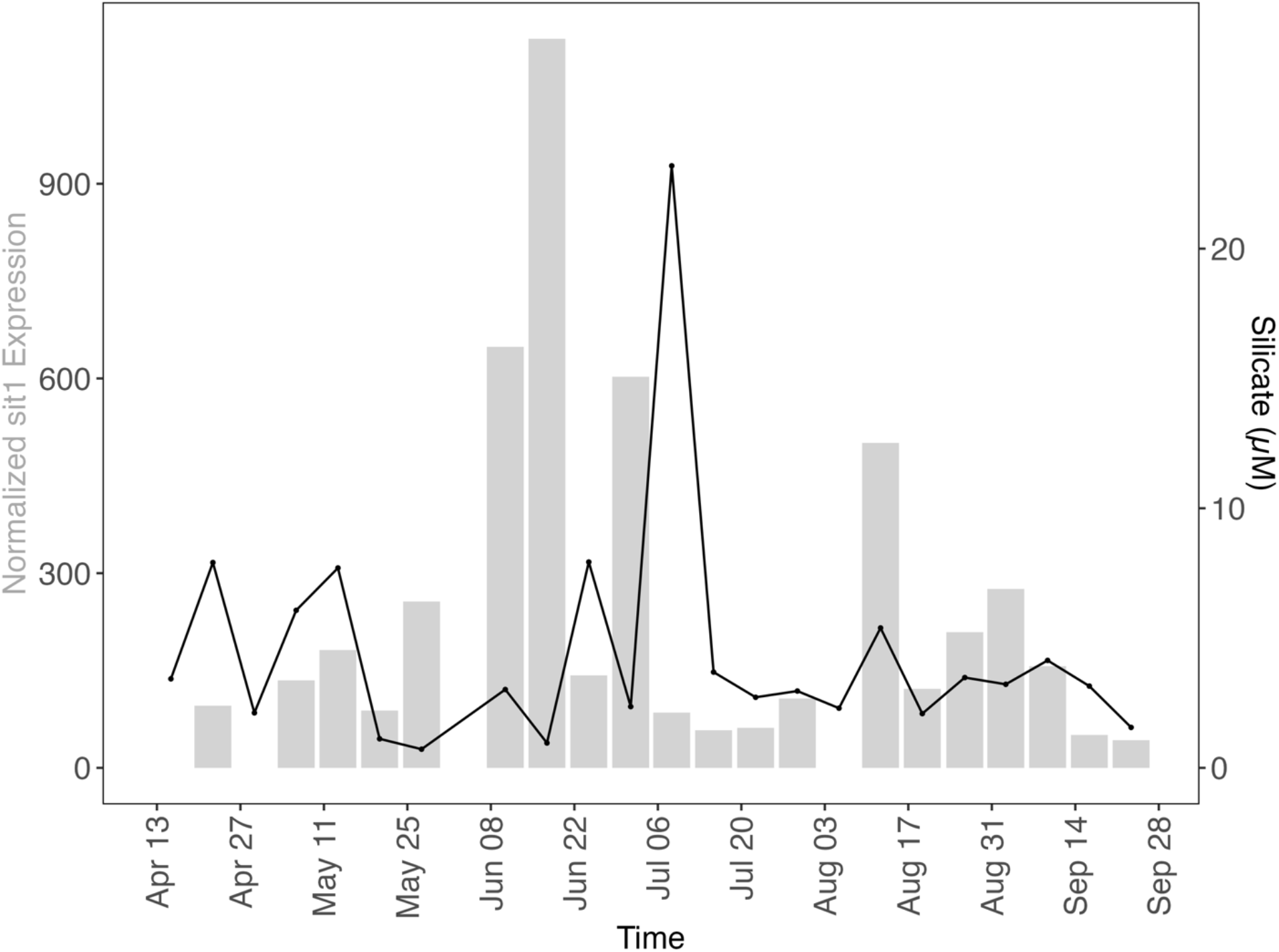
Normalized expression of *P. australis sit1* (gray bars) compared to dissolved silica concentrations (black) at MWII during the HAB. Raw read counts of *sit1* from *Pseudo-nitzschia australis* were normalized to total *Pseudo-nitzschia* read counts and multiplied by 1.0 x 10^6^.

**Figure S13.**
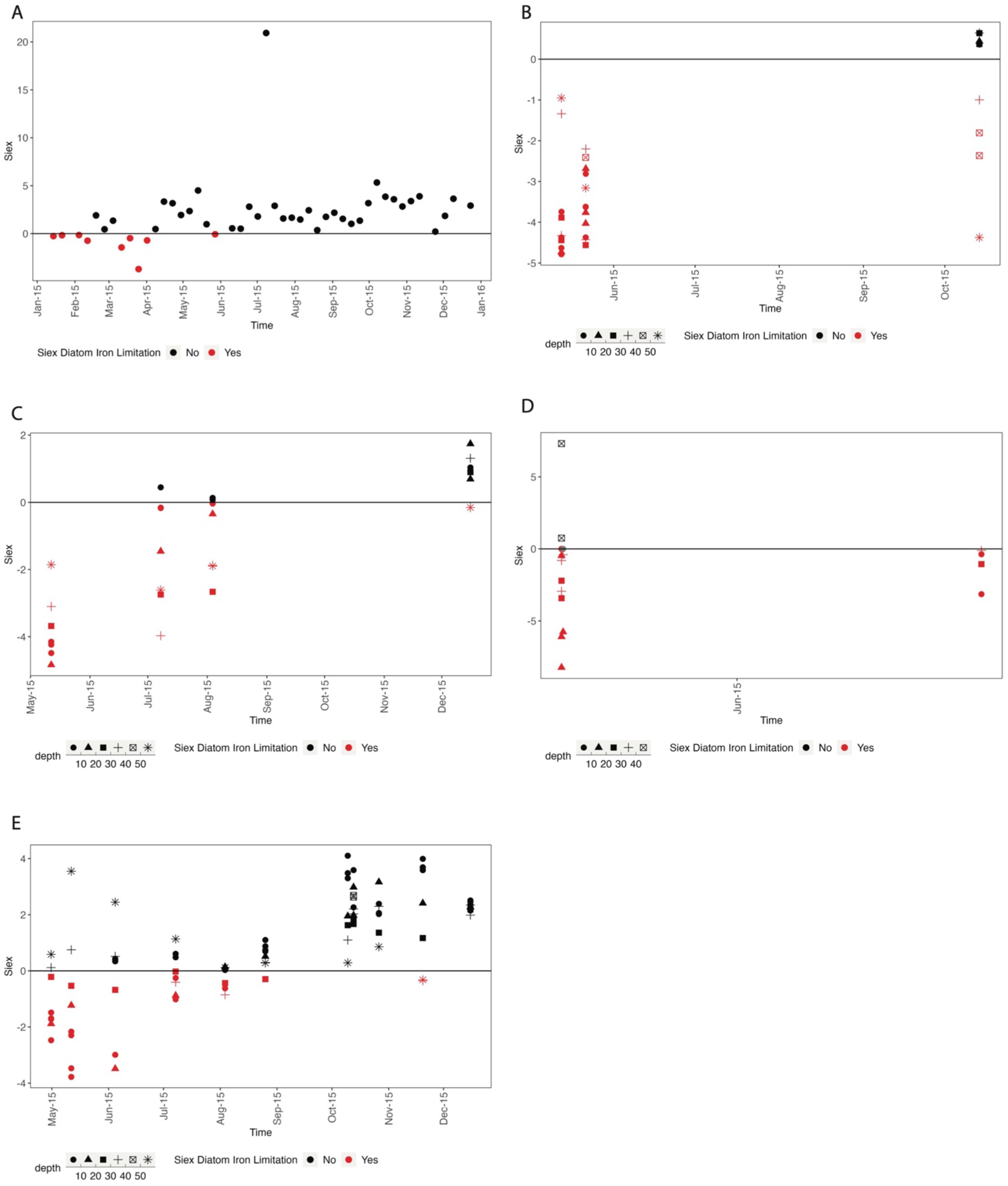
Calculations of Si_ex_, a proxy for diatom iron limitation, suggest iron limitation early in the year in Monterey Bay, CA in several locations shown in Figure S1. Negative values of Si_ex_ indicate that diatoms preferentially take up H_4_SiO_4_ relative to NO_3_^-^ due to iron deficiency, and more negative Si_ex_ values indicate a higher level of iron deficiency. All calculations assume the ratio of H_4_SiO_4_ to NO_3_^-^ at the regional upwelling source depth “R-preformed” is equal to 1, and include measurements from depths above 65 meters. Si_ex_ is calculated at (A) MWII, (B) MARS, (C) M2, (D) ESP South, and (E) C1.

**Figure S14.**
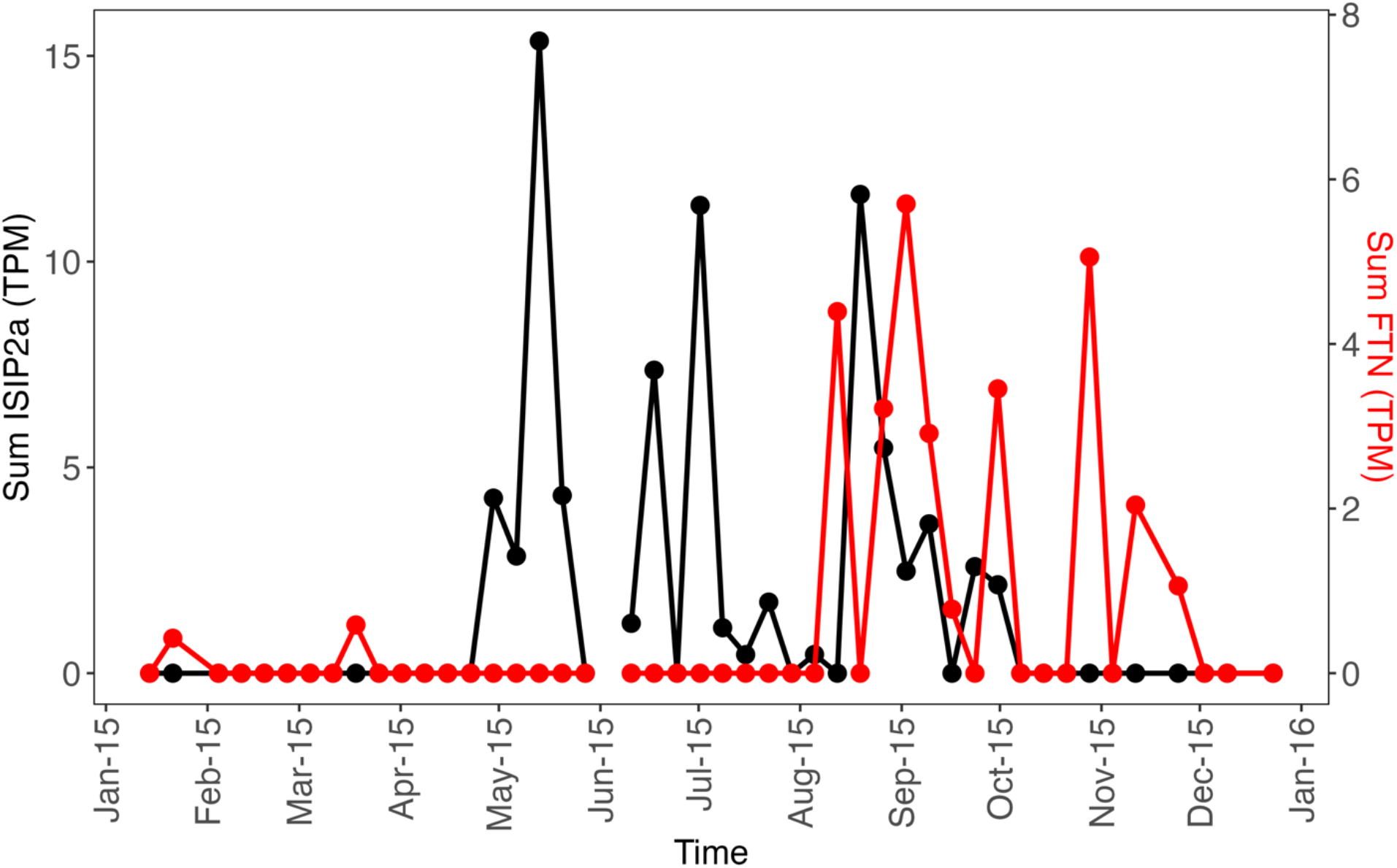
Sum of *Pseudo-nitzschia* expression of ISIP2a and ferritin (FTN) in transcripts per million (TPM).

**Figure S15.**
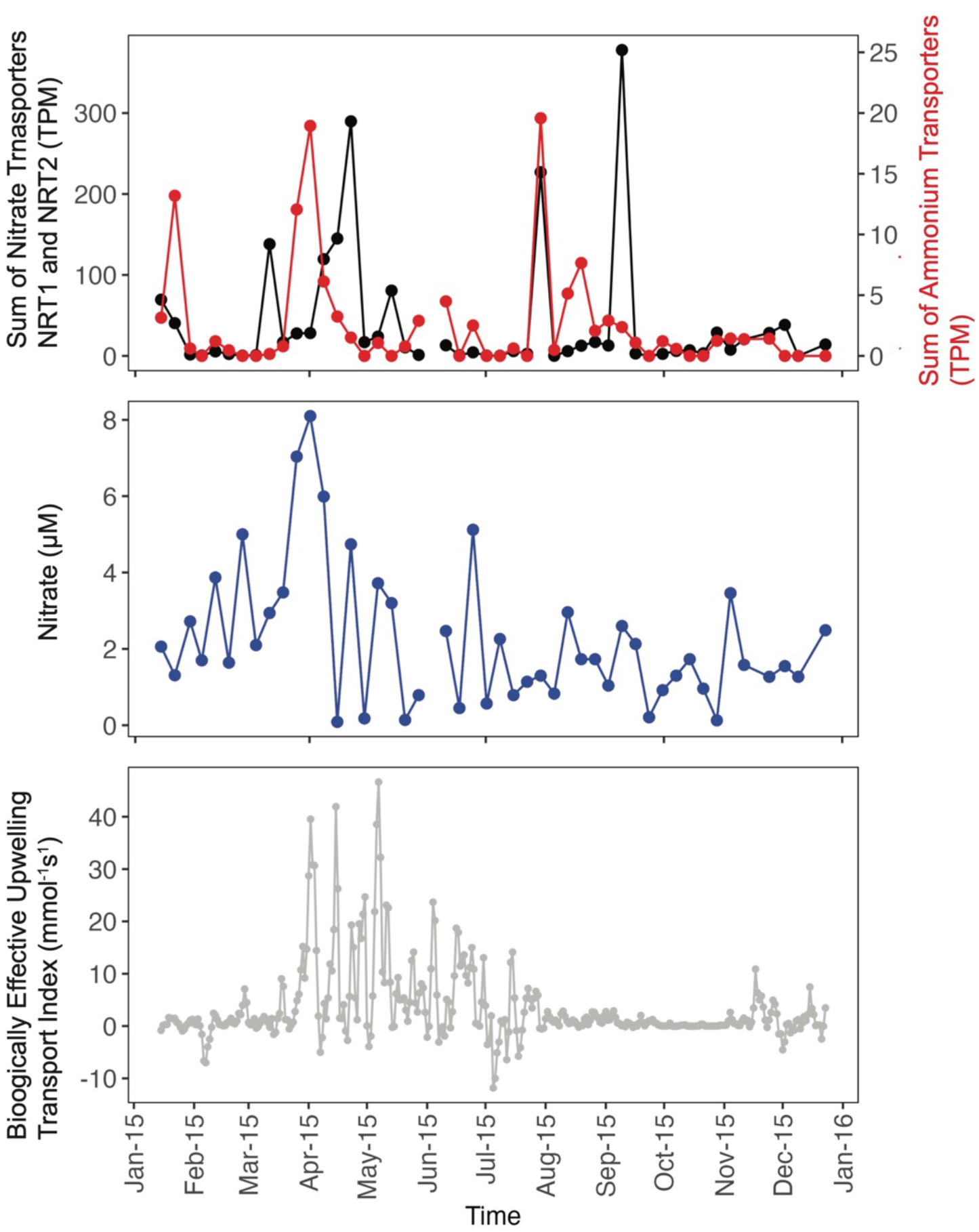
Diatom nitrogen transporters at Monterey Wharf peak in the Spring and late-Summer, reflect regional upwelling conditions.

**Figure S16.**
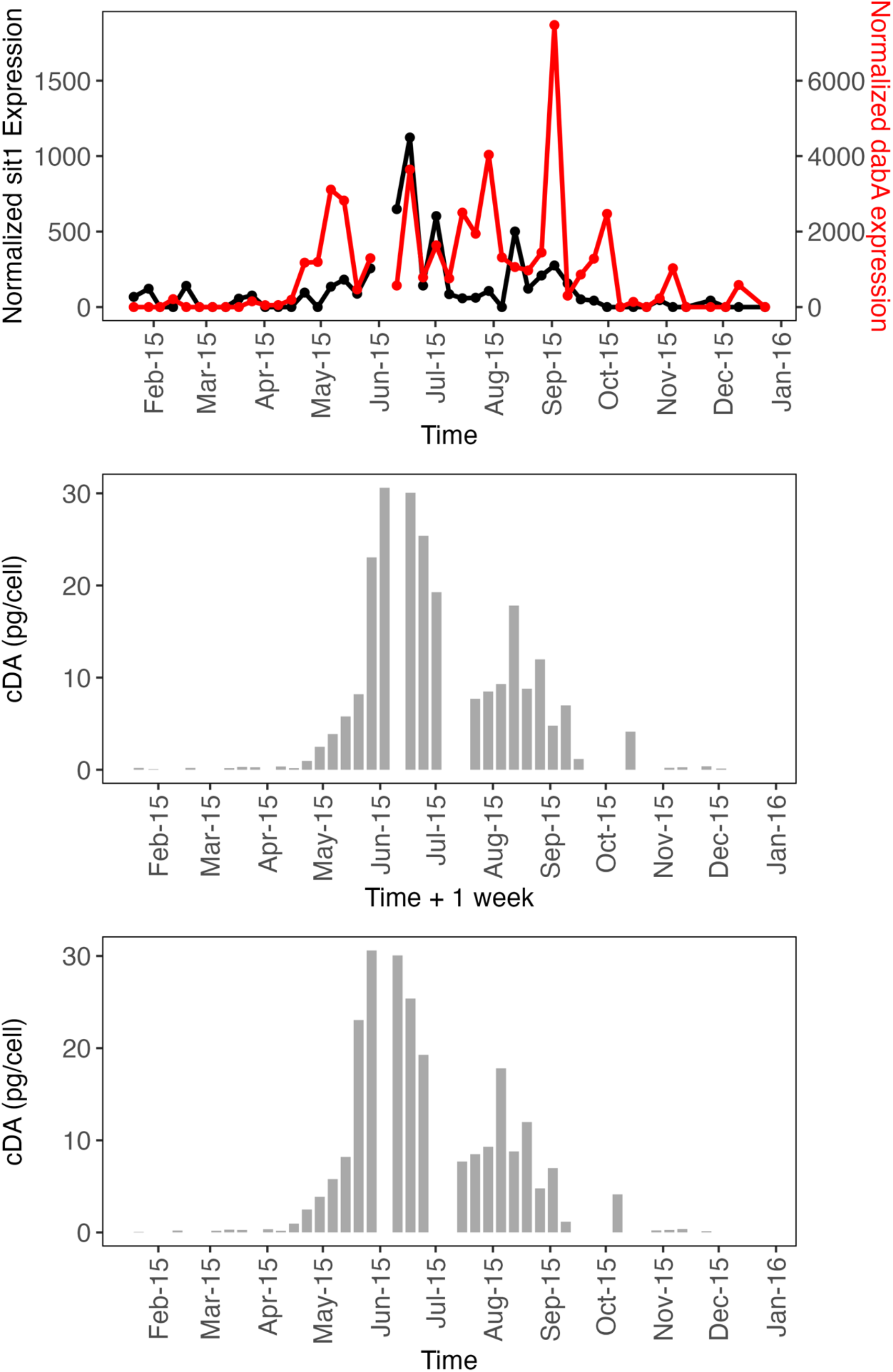
*Sit1* and *dabA* expression predict cDA in one week during the 2015 HAB. (Top) normalized *sit1* expression (black) and normalized *dabA* expression (red). Raw read counts of *sit1* and *dabA* from *Pseudo-nitzschia australis* were normalized to total *Pseudo-nitzschia* read counts and multiplied by 1.0 x 10^6^. (Middle) cDA (pg/cell) offset by one week, and (bottom) cDA (pg/cell) concurrent in time with gene expression.

**Figure S17.**
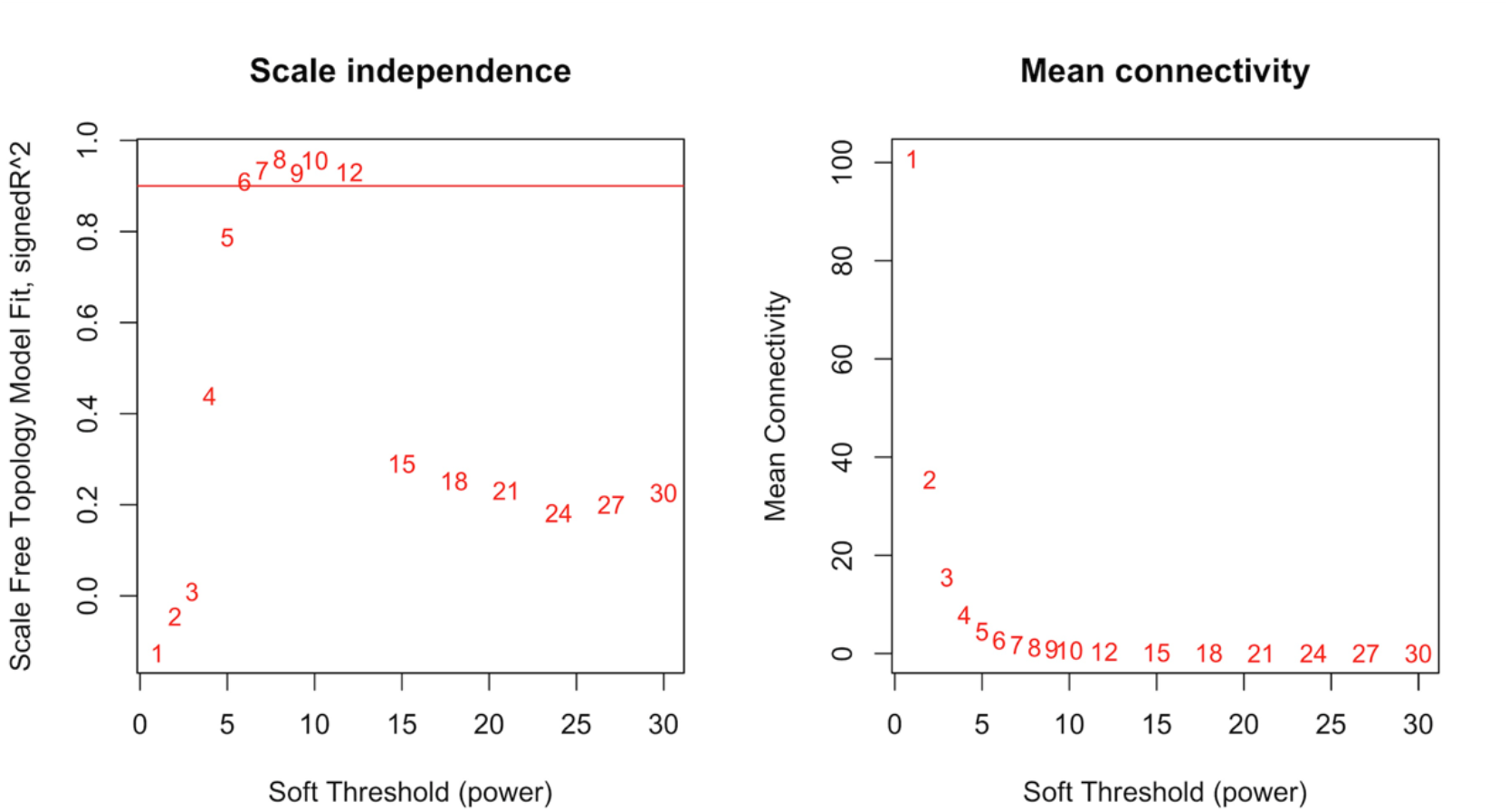
Determination of ideal soft-thresholding parameter “b” to test for the lowest value of “b” to exceed a scale-free topology R^2^ value of 0.8, showing scale-free fit index and mean connectivity.

**Figure S18.**
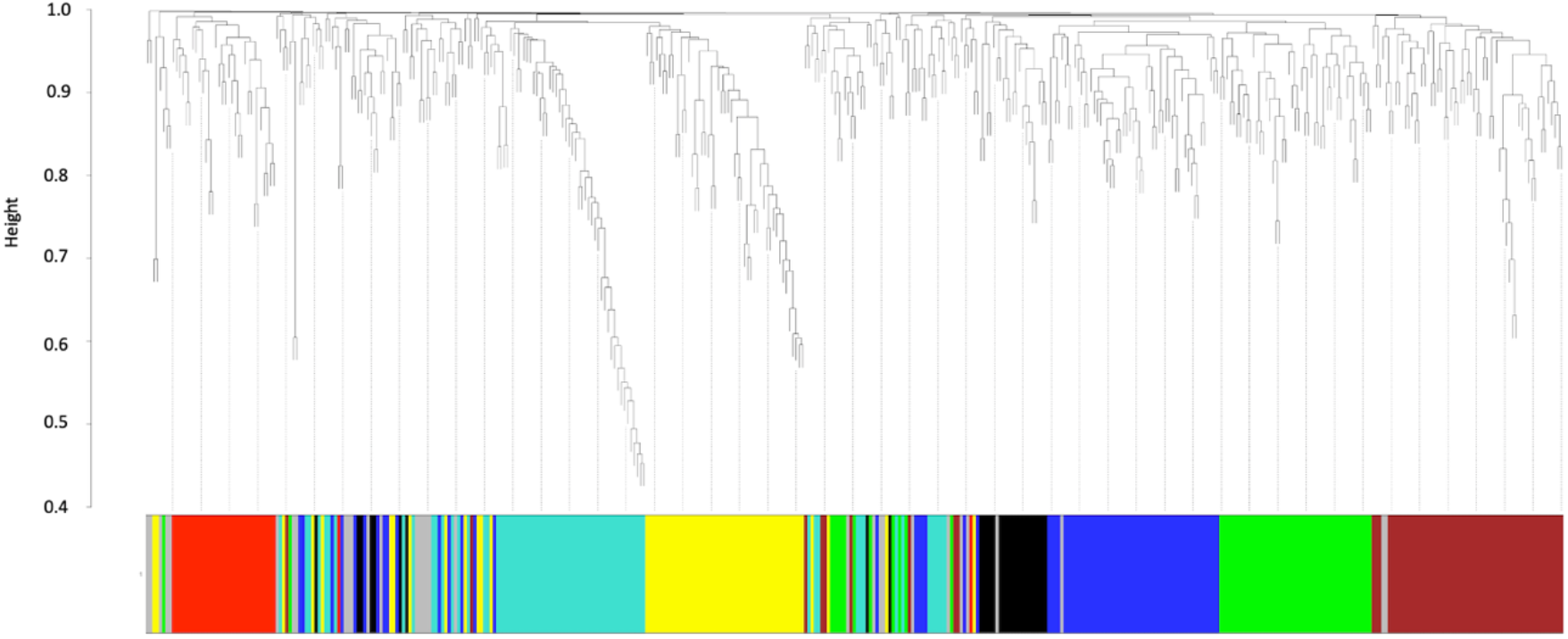
Clustering dendrogram of *P. australis* ORF expression profiles, together with assigned module colors.

**Table S1.**
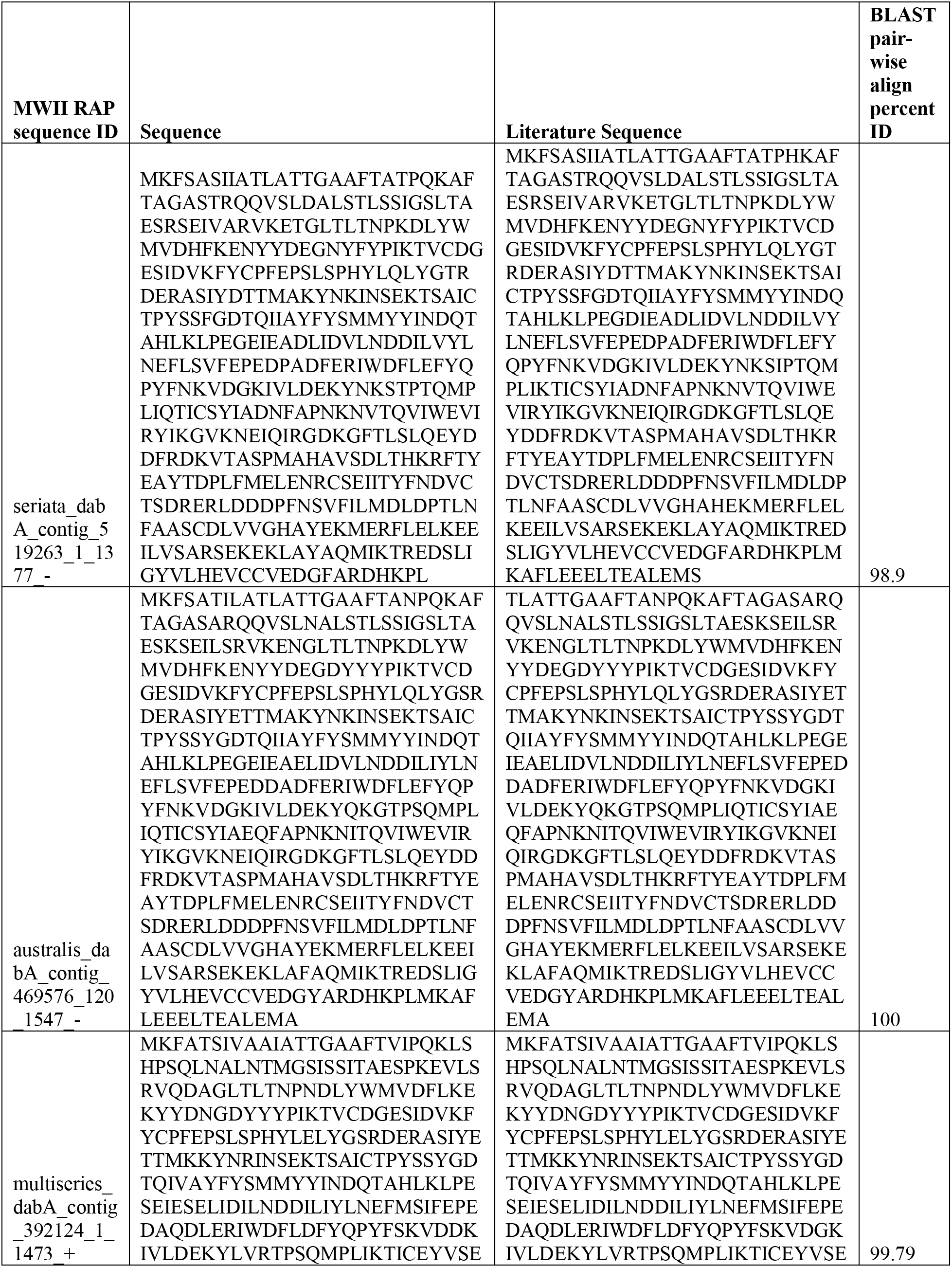
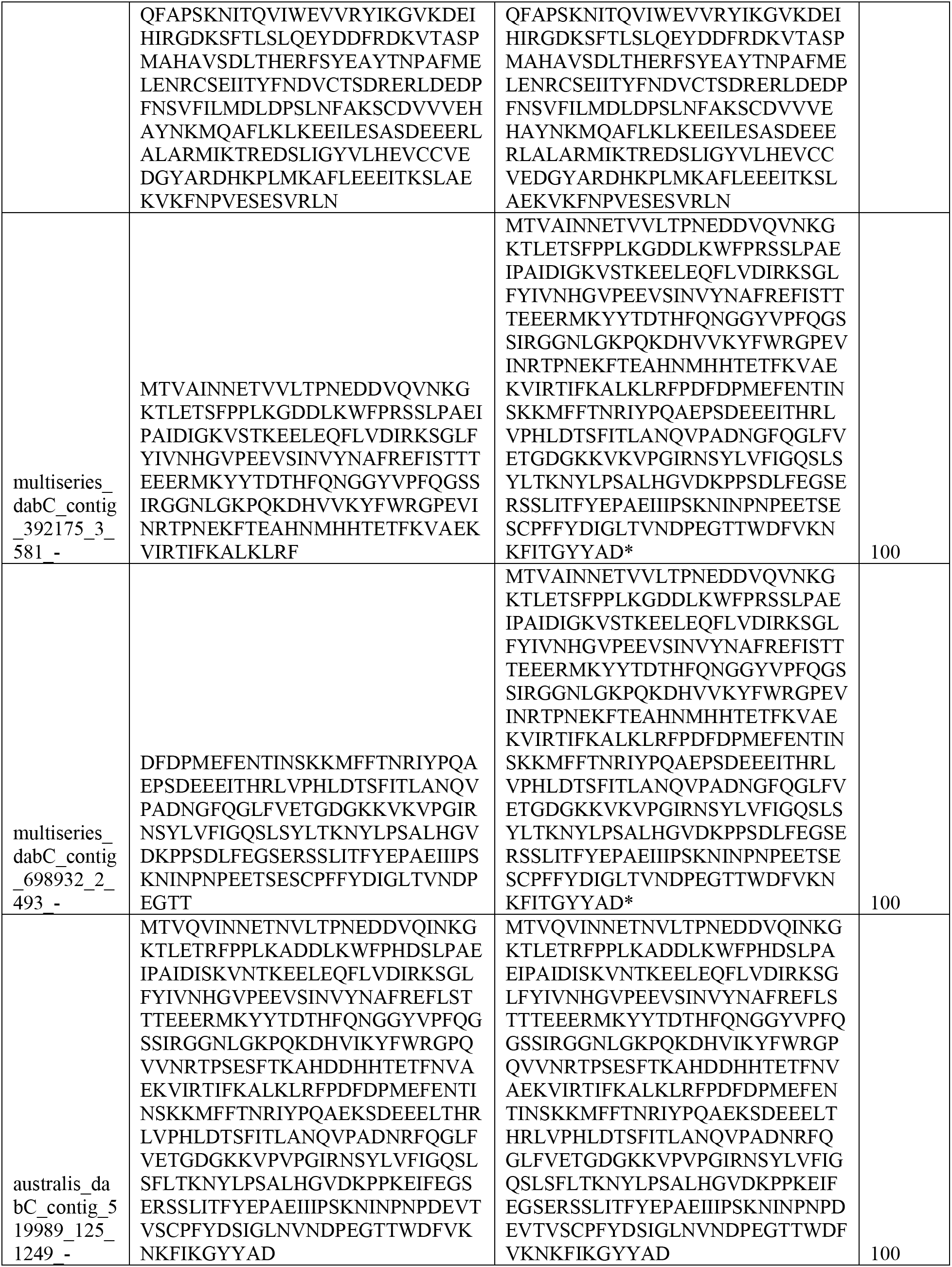
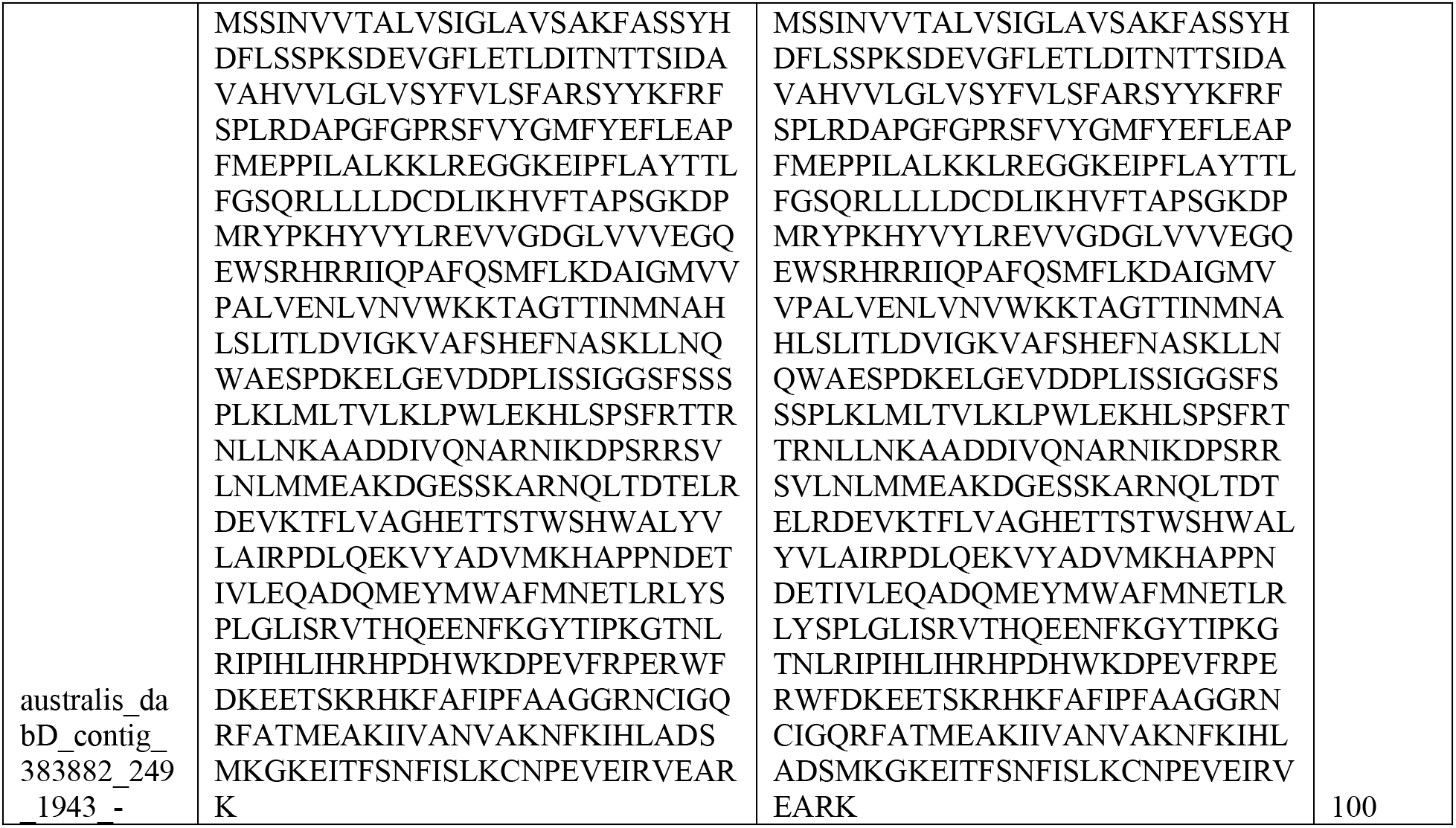
Nucleotide sequences of several *dab* transcripts in the polyA-enriched RNA-sequencing dataset compared to known *dab* genes from *P. australis, P. multiseries*, and *P. seriata*.

**Table S2.**
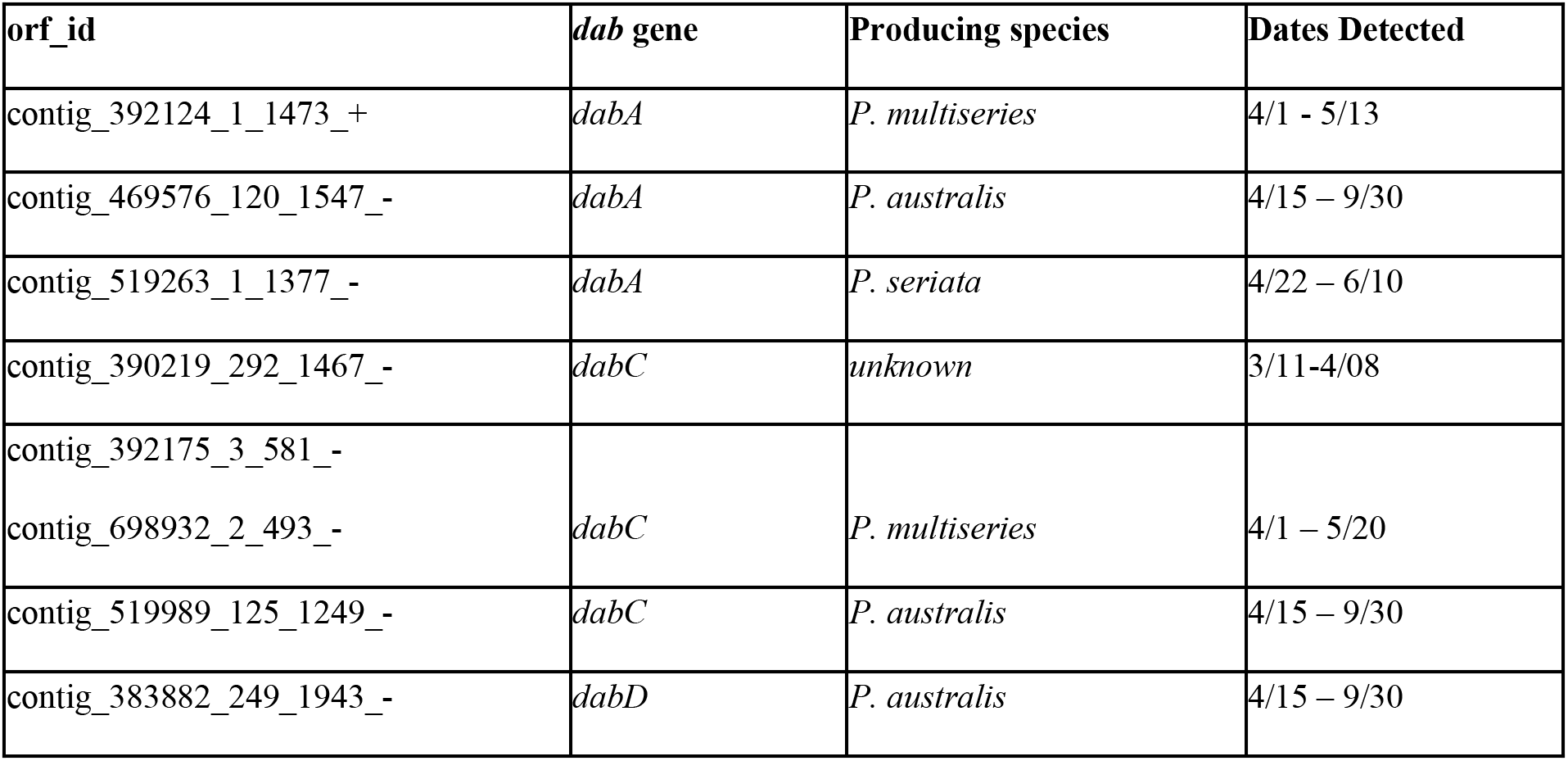
Contigs encoding *dab* genes from *de novo* metatranscriptomic assembly.

**Table S3.**
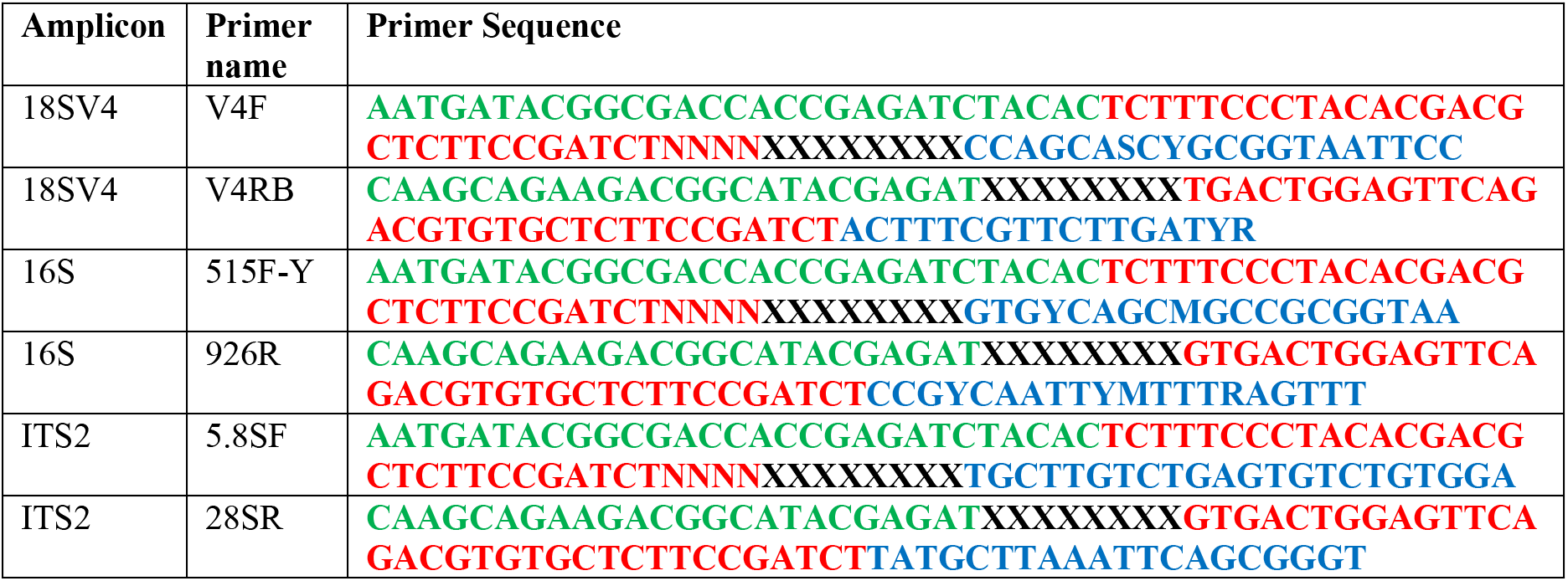
Primers used in this study. Sequence in red and green represent Illumina sequencing adaptors, black sequence represents library specific index sequences (8 nt), blue sequence represents amplicon specific annealing sequence.

**Table S4.** Domoic acid measurements from within and nearby Monterey Bay in 2015 that exceed 25 nM.

| PLATFORM | DATE | LAT | LON | DEPTH | pDA<br>(ng/L) | dDA<br>(ng/L) | dDA<br>(nM) |
| --- | --- | --- | --- | --- | --- | --- | --- |
| R/V Martin | 5/7/15 | 36.90 | -121.94 | 0 | 3051.90 | 54234.16 | 174.32 |
| R/V Martin | 5/29/15 | 36.83 | -121.85 | 0 | 240.83 | 21240.15 | 68.27 |
| R/V Martin | 5/19/15 | 36.64 | -121.92 | 12 | 12191.64 | 11817.14 | 37.98 |
| R/V Martin | 5/19/15 | 36.64 | -121.88 | 20 | 408.53 | 10607.32 | 34.09 |
| R/V Martin | 5/19/15 | 36.64 | -121.88 | 40 | 93.63 | 10189.48 | 32.75 |
| R/V Martin | 5/26/15 | 36.74 | -121.91 | 21 | 1088.85 | 8532.63 | 27.43 |
| R/V Martin | 5/29/15 | 36.83 | -121.85 | 14 | 510.48 | 8149.67 | 26.19 |
| R/V Carson | 6/5/15 | 36.61 | -122.01 | 9 | 4212.16 | 35151.18 | 112.98 |
| R/V Carson | 6/5/15 | 36.70 | -121.89 | 0 | 6361.49 | 20999.23 | 67.50 |
| R/V Carson | 6/5/15 | 36.64 | -121.88 | 0 | 4485.38 | 19881.90 | 63.90 |
| R/V Carson | 5/28/15 | 36.78 | -121.87 | 0 | 2128.14 | 17143.37 | 55.10 |
| R/V Carson | 6/5/15 | 36.78 | -121.87 | 0 | 9641.05 | 13487.89 | 43.35 |
| R/V Carson | 6/5/15 | 36.70 | -121.89 | 7 | 8301.68 | 10321.89 | 33.18 |
| R/V Carson | 5/12/15 | 36.90 | -121.93 | 0 | 807.04 | 10096.42 | 32.45 |
| R/V Carson | 5/12/15 | 36.90 | -121.93 | 8 | 1245.01 | 9312.03 | 29.93 |
| R/V Carson | 5/28/15 | 36.78 | -121.87 | 60 | 171.78 | 8901.58 | 28.61 |
| Shimada | 6/29/15 | 36.45 | -122.14 | 3 | 353.92 | 20294.34 | 65.23 |
| Shimada | 6/30/15 | 36.70 | -122.24 | 3 | 3901.48 | 20018.15 | 64.34 |
| Shimada | 6/28/15 | 36.20 | -122.15 | 3 | 1361.42 | 11981.32 | 38.51 |
| Shimada | 6/30/15 | 36.70 | -122.03 | 3 | 5040.44 | 10328.06 | 33.20 |
| Shimada | 6/29/15 | 36.45 | -122.56 | 3 | 186.00 | 8682.57 | 27.91 |
| Shimada | 6/29/15 | 36.45 | -122.34 | 3 | 223.31 | 8398.60 | 27.00 |
| Santa Cruz<br>Wharf | 11/25/15 | 36.96 | -122.02 | Mix 0,<br>1.5, and 3<br>m: equal<br>volumes | 13.35 | 980599.22 | 3151.84 |
| Santa Cruz<br>Wharf | 7/31/15 | 36.96 | -122.02 | Mix 0,<br>1.5, and 3<br>m: equal<br>volumes | 132.77 | 12427.17 | 39.94 |

**Table S5.**
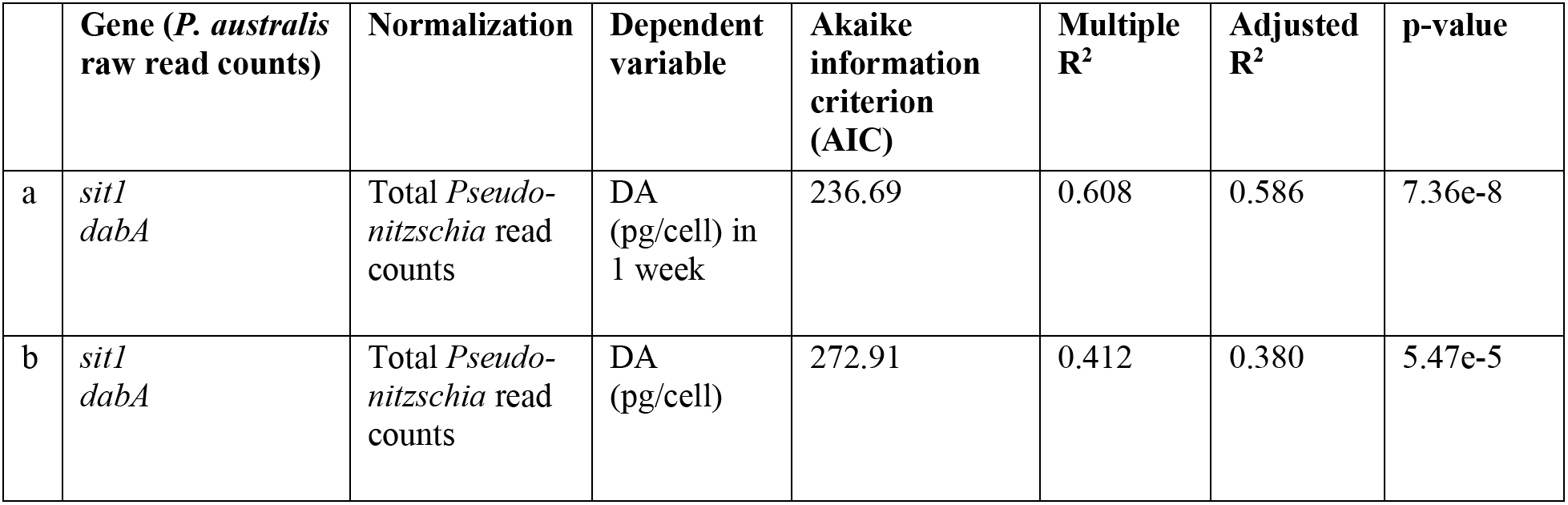
Statistical results from multivariable generalized linear models used to predict domoic acid. Independent variable gene expression was specifically from *Pseudo-nitzschia australis*, and all raw counts were normalized to total *Pseudo-nitzschia* read counts.

